# TaNF-Y-PRC2 orchestrates temporal control of starch and protein synthesis in wheat

**DOI:** 10.1101/2023.12.26.573020

**Authors:** Jinchao Chen, Long Zhao, Haoran Li, Changfeng Yang, Dongzhi Wang, Xuelei Lin, Yujing Lin, Hao Zhang, Xiaomin Bie, Peng Zhao, Shengbao Xu, Xiansheng Zhang, Xueyong Zhang, Yingyin Yao, Jun Xiao

## Abstract

The endosperm in cereal grains is instrumental in determining grain yield and seed quality, as it controls the production of starch and protein. In this study, we identified a specific TaNF-Y trimeric complex, consisting of TaNF-YA3-D, TaNF-YB7-B, and TaNF-YC6-B, exhibiting robust expression within endosperm during grain filling stage in wheat. Knock-down of either *TaNF-YA3* or *TaNF-YC6* led to less starch but more gluten proteins. Detailed analyses have unveiled that the TaNF-Y indirectly boosts starch biosynthesis genes by reducing TaNAC019, a repressor of *TaAGPS1a, TaSuS2*, thereby regulating starch biosynthesis. Conversely, the TaNF-Y directly inhibits the expression of gliadin and low molecular weight (LMW)-GS coding genes, including *TaGli-γ-700* and *TaLMW-400*. Furthermore, the TaNF-Y components interact with TaSWN, the histone methyltransferase subunit of Polycomb repressive complex 2 (PRC2), to repress the expression of *TaNAC019*, *TaGli-γ-700*, and *TaLMW-400* through H3K27me3 modification. Notably, weak mutation of *TaFIE*, core subunit of PRC2, has reduced starch but elevated gliadin and LMW-GS levels. Intriguingly, DNA variations of TaNF-Y components are widely associated with seed developmental traits. In particular, variation within the coding region of *TaNF-YB7-B* is linked to differences in starch and protein content. Distinct haplotypes of *TaNF-YB7-B* affect its interaction with TaSWN, influencing the repression of targets like *TaNAC019* and *TaGli-γ-700*. Our findings illuminate the intricate molecular mechanisms governing epigenetic regulation by the TaNF-Y-PRC2 for wheat endosperm development. Manipulating the TaNF-Y complex holds potential for optimizing grain yield and enhancing quality.

## Introduction

Wheat is one of the most important staple crops worldwide, providing a significant portion of the global population’s dietary needs (Shewry and Hey, 2015). Ensuring high grain yield and superior quality in wheat is crucial for food security (Xiao et al., 2022). Grain yield, determined by the accumulation of starch in the endosperm, is a primary target for crop improvement. Starch biosynthesis, a complex metabolic pathway, involves several key enzymes, such as ADP-glucose pyrophosphorylase (AGPase), granule-bound starch synthases (GBSS), starch synthases (SSs), and starch branching enzymes (SBEs), which together orchestrate the intricate process of starch production (Pfister and Zeeman, 2016; Liu et al., 2013; Ohdan et al., 2005; Han et al., 2007). Simultaneously, starch-degrading enzymes like starch phosphorylase (PHO) and amylases play a pivotal role in determining starch content and quality during grain filling (Cuesta-Seijo et al., 2017; Subasinghe et al., 2014).

Numerous transcription factors (TFs) that regulate starch biosynthesis in wheat have been identified, such as TubZIP28/TabZIP28, TaRSR1, and TaNAC019 (Xiao et al., 2022). Knockdown of TabZIP28 leads to reduction in the expression and activity of cytoplasmic AGPases, resulting in approximately 4% decrease of total starch content within mature kernels (Song et al., 2020). Additionally, TaRSR1, an essential transcriptional repressor, temporally regulate the expression of starch biosynthesis-related enzymes (Liu et al., 2016). Overexpression of *TaNAC019* downregulated grain weight and starch content (Liu et al., 2020). Whereas, another study showed that *TaNAC019* positively regulates genes involved in starch synthesis and storage protein accumulation (Gao et al., 2021). Therefore, the exact role of TaNAC019 in wheat starch synthesis and glutenin accumulation remains unclear.

In parallel, end-use quality, with a particular focus on gluten protein content and composition, assumes paramount significance for the nutritional value and end-use applications of wheat. Gluten proteins, comprising gliadins and glutenins, exert profound influence over the dough quality and end-use characteristics of wheat flour (Biesiekierski, 2017). Regulating gluten protein accumulation involves a complex interplay among a network of genes and transcription factors. Key regulators, such as high molecular weight glutenin subunit (HMW-GS), low molecular weight glutenin subunit (LMW-GS), and gliadin, act as drivers of gluten protein gene expression in the endosperm, with varying ratios of these proteins being essential for different dough applications (Dong et al., 2010; Liu et al., 2005; Payne et al., 1987; Shewry et al., 2002; Rasheed et al., 2014; Veraverbeke and Delcour, 2002). Additionally, transcription factors like Dof (DNA-binding with one finger) proteins and bZIP (basic leucine zipper) factors contribute to the modulation of gluten protein synthesis, ensuring precise spatiotemporal control over gluten protein accumulation during seed development.

The NF-Y (nuclear factor Y) complex, composed of NF-YA, NF-YB, and NF-YC subunits, is a conserved transcription factor that plays a crucial role in seed development in different plant species (Petroni et al., 2012). This trimeric complex binds to the CCAAT-box cis-element in the promoters of target genes (Gnesutta et al., 2017; Laloum et al., 2013; Nardone et al., 2017). The NF-Y complex has been well-documented in influencing embryo development, seed filling, starch synthesis, and accumulation in rice (Feng et al., 2022; Xu et al., 2016; Bello et al., 2019). In maize, the NF-Y complex regulates endosperm development, impacting grain size and weight (Zhang et al., 2022). While the specific role of the NF-Y complex in wheat seed development is still under investigation, preliminary studies suggest its involvement in regulating grain filling and seed quality (Liu et al., 2023; Yadav et al., 2015). In addition, how NF-Y complex regulates starch and/or storage protein coordinately in cereals is unclear.

Epigenetic regulation, encompassing histone modifications, emerges as a crucial factor in controlling seed development, particularly during the early stages of endosperm development (Ding et al., 2022). Among these modifications, the Polycomb Repressive Complex 2 (PRC2) and its catalyzed mark, H3K27me3, stand out as pivotal players. PRC2 and H3K27me3 dynamically regulate gene expression during the initiation of endosperm proliferation and cellularization, ensuring proper development and contributing to grain filling and yield. This regulatory mechanism has been extensively studied in various plant species, such as rice, maize, wheat, and Arabidopsis (Cheng et al., 2020,2021; Tonosaki et al., 2021; Ni et al., 2019; Zhang et al., 2018; Gutierrez-Marcos et al., 2013; Zhang et al., 2023). However, the function of PRC2 and H3K27me3 in regulating later endosperm development processes, such as starch biosynthesis and gluten protein content, remains largely uncharted territory, particularly in wheat.

Here, we investigated the role of an endosperm-expressed TaNF-Y complex in regulating starch biosynthesis and gluten protein accumulation in wheat. Our results reveal that this regulation involves TaNF-Y mediated recruitment of PRC2 and direct H3K27me3 deposition on gluten protein coding genes and indirect modulation of starch biosynthesis through repression of *TaNAC019*. Additionally, natural variations in the *TaNF-YB7-B* component impact starch and protein content by influencing interaction with PRC2 and alteration of expression of targets.

## Results

### Presence of endosperm highly expressed TaNF-Y in wheat

During endosperm development, starch and seed storage protein (SSP) biosynthesis are temporally regulated (Wei et al., 2009, Yin et al., 2012). Through a time-serial transcriptome analysis (Zhao et al., 2023), we observed an earlier expression of starch biosynthesis related genes, primarily peaking at 6-8 DAP (days after pollination), in contrast to the relatively later and broader expression window of various SSPs spanning from 6 to 16 DAP (Figure 1A-B and Supplemental Data Set S1). We wonder how such temporal expression pattern is orchestrated by various TFs. Notably, known regulators such as TaNAC019 and TaPBF exhibit varied dynamic expression patterns throughout the entire developmental period from 0 DAP to 22 DAP (Figure 1B). Furthermore, we found a component of the TaNF-Y complex, TaNF-YA3, exhibits a transcriptional profile similar to that of starch biosynthesis genes, as evidenced by the transcriptome data (Figure 1B).

**Figure 1.**
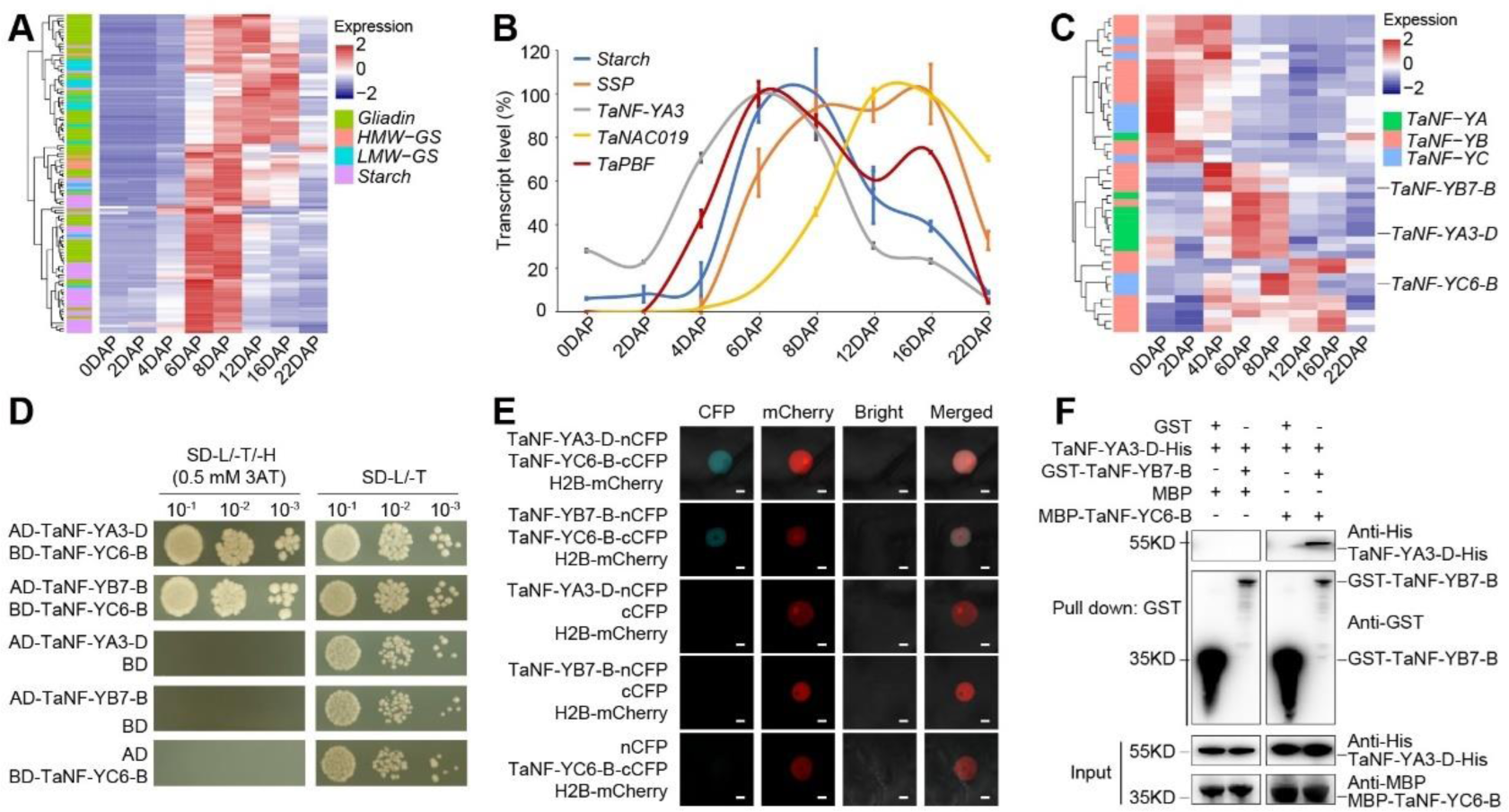
The endosperm-expressed trimeric TaNF-Y complex in wheat. A. The expression heatmap of starch and SSP related genes during endosperm development in wheat. DAP, days after pollination. B. The percentile transcript level of *TaNF-YA3, TaPBF, TaNAC019* and starch synthesis gene *AGPS1a*, and SSP synthesis gene *TaGli-γ-700* during the course of endosperm development. The relative expression of each gene compared to *TaActin* was calculated. Maximum expression of each gene is set as 100%, and relative levels are shown. Reported as mean ± SD of three biological replicates. C. The expression heatmap of endosperm expressed TaNF-Y components genes during endosperm development in wheat. D. Y2H assay showing the interaction between TaNF-YA3-D and TaNF-YC6-B, TaNF-YB7-B and TaNF-YC6-B. SD-L/-T/-H, (SD-Leu/-Trp/-His, 0.5mM 3AT); SD-L/-T, (SD-Leu/-Trp). E. BiFC assay of the interaction between TaNF-YA3-D and TaNF-YC6-B, TaNF-YB7-B and TaNF-YC6-B in tobacco leaves. H2B-mCherry was used as cell nucleus marker. Scale bar, 0.5 µm. F. Analysis of the interaction of TaNF-YA3-D-His, MBP-TaNF-YC6-B with GST-TaNF-YB7-B recombinant proteins with a GST pull-down assay.

In hexaploid wheat, a total of 19 NF-YA, 36 NF-YB, and 18 NF-YC subunits have been identified through sequence similarity alignment (Supplemental Figure S1A). These coding triads generally exhibit similar transcriptional patterns across different tissues, showcasing balanced expression levels (Supplemental Figure S1B-C). Notably, certain TaNF-Y coding genes demonstrate high expression during grain and spike development (Supplemental Figure S1B). Among these, *TaNF-YA3-D* (*TraesCS4D02G289600*), *TaNF-YB7-B* (*TraesCS7B02G121900*), and *TaNF-YC6-B* (*TraesCS6B02G185700*) display specifical expression during the 4-12 DAP period of seed development (Figure 1C), with no reported functional information regarding their orthologues in other cereals (Xu et al., 2016; Zhang et al., 2022). The interaction between TaNF-YA3-D/TaNF-YC6-B, and TaNF-YB7-B/TaNF-YC6-B within nuclei has been confirmed through yeast two-hybridization (Y2H) and bi-fluorescence complementation (BiFC) assays (Figure 1D-E), aligning with their subcellular localization (Supplemental Figure S1D). Consistent with findings in other species (Hou et al., 2014), no direct interaction between TaNF-YA3-D and TaNF-YB7-B was observed (Figure 1F). Remarkably, the addition of TaNF-YC6-B facilitated the interaction between TaNF-YA3-D and TaNF-YB7-B via an *in vitro* pull-down assay (Figure 1F), suggesting their potential to form a trimeric complex. Therefore, we have identified a TaNF-Y complex dynamically expressed in the endosperm of hexaploid wheat.

### TaNF-Y affects starch synthesis and gluten protein composition

During 6-12 DAPs, wheat grain filling begins, characterized by the synthesis of starch and storage proteins (Zhang et al., 2021). Given the spatiotemporal expression pattern of the TaNF-Y in the endosperm during this stage, we speculate its involvement in the regulation of starch and protein biosynthesis. To investigate this, we generated multiple knock-down lines of *TaNF-YA3* and *TaNF-YC6* with varying dosages using an RNAi strategy (Supplemental Figure S2A-C) (Borrill et al., 2015; Sestili et al., 2019). In the T2 generations, we observed a significant reduction in grain length (GL), grain width (GW), and thousands of grain weight (TGW) in multiple *TaNF-YA3 RNAi* and *TaNF-YC6 RNAi* lines (Figure 2A-C). Since starch serves as the major storage material in seeds (Toepfer et al., 1972), we further quantified starch content. Consistent with the smaller seed size, both total starch, amylose content was significantly reduced in *TaNF-YA3 RNAi* and *TaNF-YC6 RNAi* lines (Figure 2D, E). Based on coomassie brilliant blue staining of semi-thin sections, the *TaNF-YA3-RNAi* and *TaNF-YC6-RNAi* lines contained less starch granules (SG) compared with the Fielder (Figure 2F). Scanning electron microscopy (SEM) (Liu et al., 2020) revealed that smaller size of A-Type SGs and reduced number of B-Type SGs in *TaNF-YA3 RNAi* and *TaNF-YC6 RNAi* lines as compared to Fielder (Supplemental Figure S3A-C).

**Figure 2.**
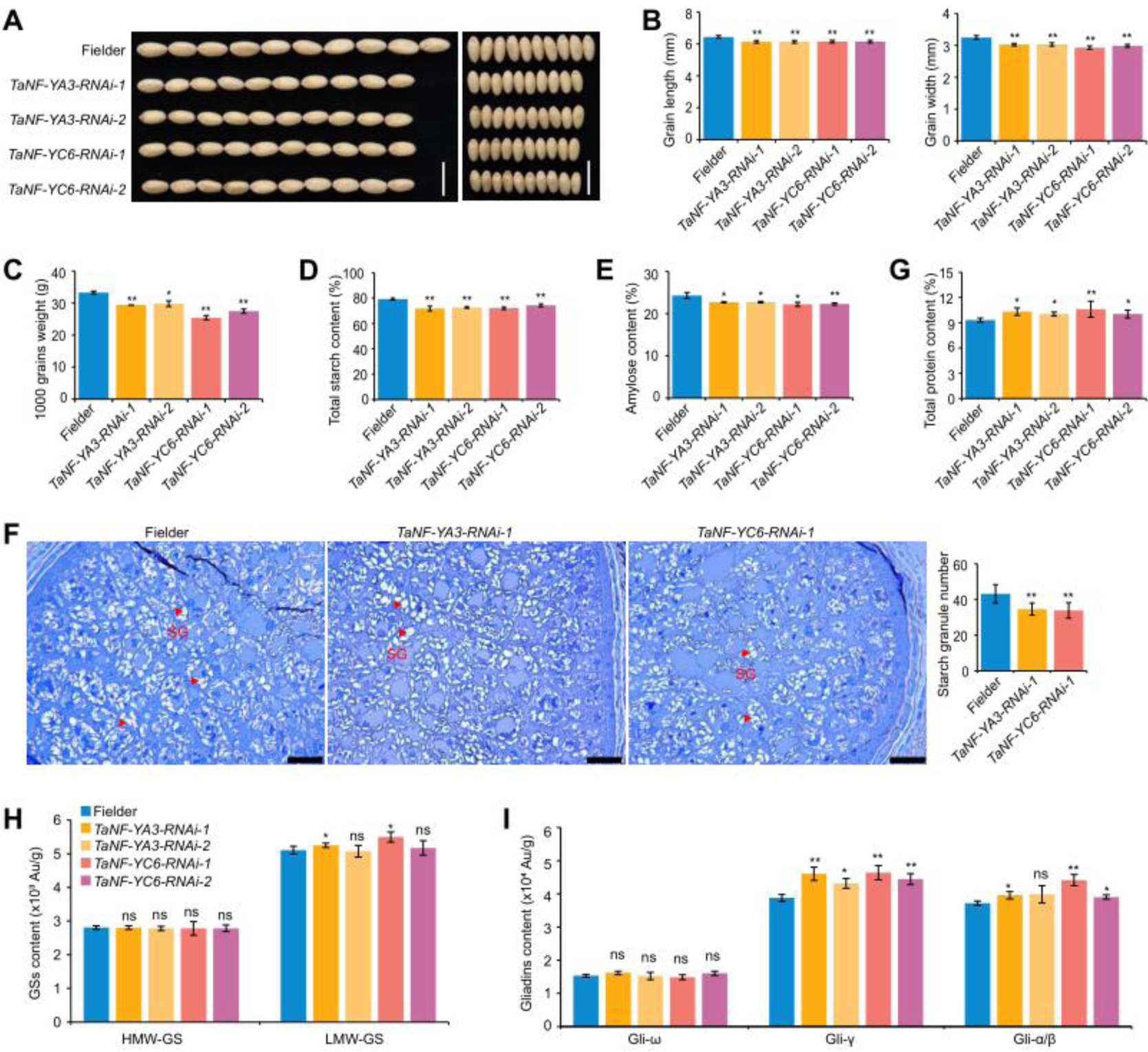
Knock-down of *TaNF-YA3* and *TaNF-YC6* influence starch biosynthesis and SSP composition. A. Grain morphology of knock-down lines for *TaNF-YA3* and *TaNF-YC6*, and transgenic null lines (Fielder). Scale bars, 1 cm. B - F. Statistical comparisons of grain length (GL) and grain width (GW) (B), thousand grain weight (TGW) (C), total starch content (D), amylose content (E), Starch granules analysis of *TaNF-YA3-RNAi-1*, *TaNF-YC6-RNAi-1* and Fielder (F), The 10 DAP seeds were fixed in FAA, and then cut into semi-thin sections after embedding with resin, stained with coomassie brilliant blue, and observed. The red triangles represent starch granules (SG). Scale bar, 100 µm. Statistical significance was determined by Student’s *t* test. *, *P* < 0.05. H-I. Total protein content (H), and the result from RP-HPLC analysis of HMW-GS, LMW-GS (I), gliadins(J) contents. Grain size-related traits including TGW, grain length and width were determined using a camera-assisted phenotyping system with five biological replicates, each with approximately 10 g seeds. The contents of total protein, HMW-GS, LMW-GS were estimated by RP-HPLC analysis with 3 biological replicates. Statistical significance was determined by Student’s *t* test. *, *P* < 0.05; **, *P* < 0.01; ns, no significant difference.

Moreover, the knock-down of *TaNF-YA3* and *TaNF-YC6* function was associated with an increase in total protein contents (Figure 2G). We also performed reverse-phase high-performance liquid chromatography (RP-HPLC) to measure HMW-GS and LMW-GS contents in *TaNF-Y* knock-down lines. The levels of HMW-GS were similar in the *TaNF-YA3-RNAi* and *TaNF-YC6-RNAi* lines as compared to Fielder (Figure 2H). Whereas, the LMW-GS levels were only increased in one line of *TaNF-YA3-RNAi* and *TaNF-YC6-RNAi* with more reduction dosage (Supplemental Figure S2C), respectively (Figure 2H). However, the gliadin levels of subunits of Gli-γ and Gli-α/β were increased in multiple *TaNF-YA3-RNAi* and *TaNF-YC6-RNAi* lines compared to Fielder (Figure 2I). Thus, the TaNF-Y is involved in the regulation of starch synthesis and gluten protein composition during endosperm development.

### TaNF-Y directly represses the transcription of gluten coding genes

To address how TaNF-Y affects starch biosynthesis and gluten protein composition, we performed transcriptome analysis using the *TaNF-YA3 RNAi-1* and *TaNF-YC6 RNAi-1* lines compared to Fielder at 10 DAP, with three biological replicates (Supplemental Figure S4A). We identified 5,714 and 3,989 differentially expressed genes (DEGs) in the *TaNF-YA3 RNAi-1* and *TaNF-YC6 RNAi-1* lines, respectively, compared to Fielder (Supplemental Figure S4B and Supplemental Data Set S2). Notably, a significant overlap of DEGs emerged, indicating the functional interplay of TaNF-YA3 and TaNF-YC6 (Figure 3A). We focused on the overlapped DEGs for further analysis (Supplemental Data Set S2). Among them, 988 genes were up-regulated. Given the increased levels of gliadins and LMW-GS proteins in *TaNF-YA3* and *TaNF-YC6* knock-down lines, we investigated storage protein coding genes, including gliadins and HMW-GS and LMW-GS, which exhibited increased expression in both *TaNF-YA3 RNAi-1* and *TaNF-YC6 RNAi-1* lines (Figure 3B). This was further validated individually by qPCR across multiple *TaNF-YA3* and *TaNF-YC6 RNAi* lines (Figure 3D). Conversely, the 970 down-regulated genes encompassed *TaAGPS1a*, *TaAGPL*, and *TaGBSSII*, with a notable enrichment in GO terms associated with starch biosynthetic and metabolic process (Figure 3A, C and Supplemental Figure S4C). qPCR validation confirmed the down-regulation of these genes in both *TaNF-YA3* and *TaNF-YC6 RNAi* lines (Figure 3D).

**Figure 3.**
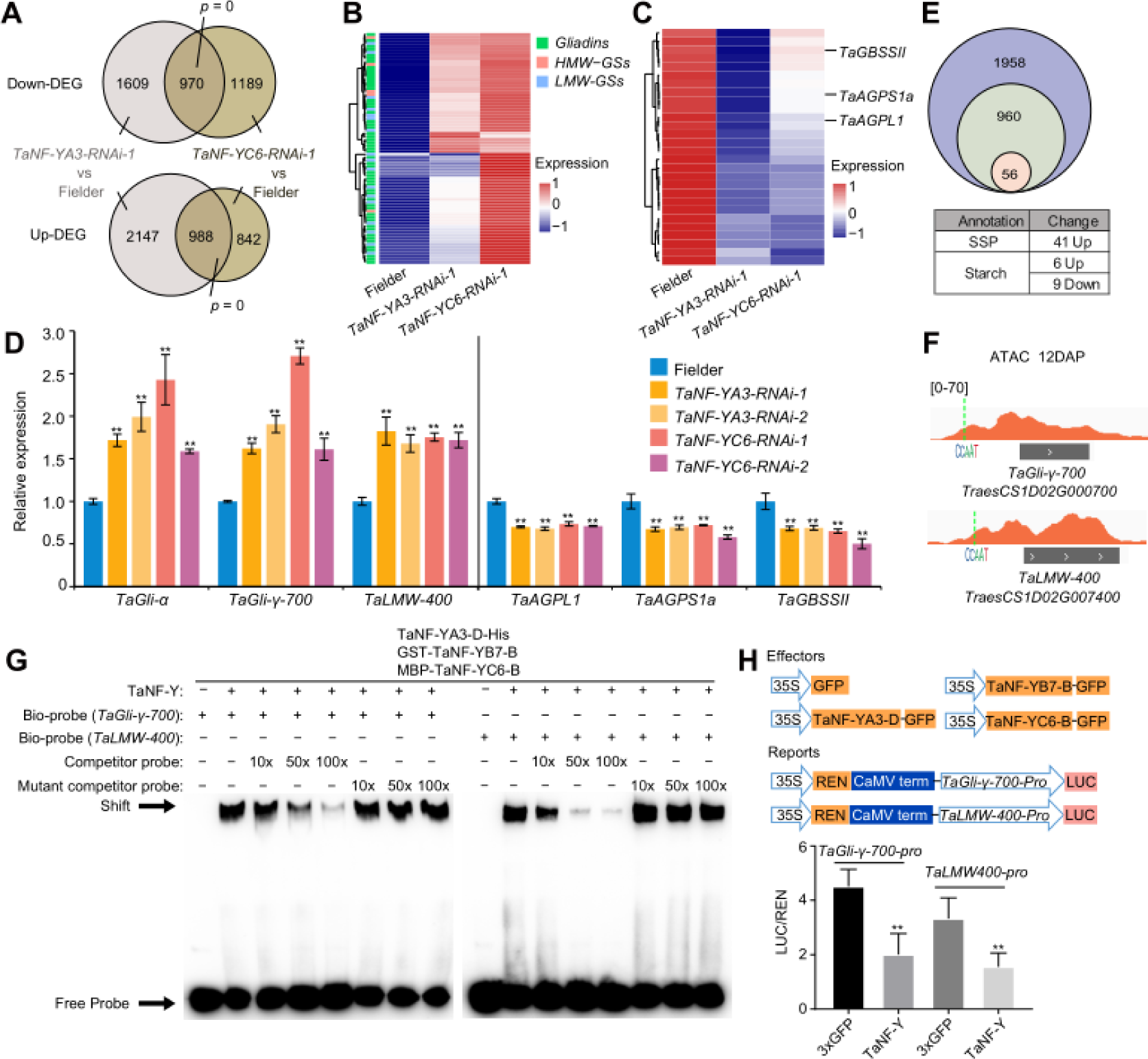
TaNF-Y directly represses the transcription of gluten coding genes to regulate SSP composition. A. The Venn-diagram shows the overlapped of up-regulated and down-regulated genes in *TaNF-YA3* and T*aNF-YC6* knock-down lines as compared to Fielder. B. Heatmap showing the expression level of SSP synthesis genes in *TaNF-YA3-RNAi-1, TaNF-YC6-RNAi-1* and Fielder. C. Heatmap showing the expression level of starch biosynthetic genes in *TaNF-YA3-RNAi-1, TaNF-YC6-RNAi-1* and Fielder. D. The relative expression level of *TaGli-α*, *TaGli-γ-700*, *TaLMW-400*, *TaAGPL1*, *TaAGPS1a* and *TaGBSSII* in the *TaNF-YA3* and *TaNF-YC6* knock-down lines, estimated by qRT-PCR and normalized to *TaActin*. The relative expression level was reported as mean ± SD of three biological replicates. Statistical significance was determined by Student’s *t* test. *, *P* < 0.05; **, *P* < 0.01. E. The diagram shows the overlap of DEGs regulated by *TaNF-YA3* and *TaNF-YC6* (identified by RNA-seq) and TaNF-Y potential target genes (identified by ATAC-seq). F. IGV shows the ATAC-seq peak of *TaGli-γ-700* and *TaLMW-400*. The CCAAT motif was indicated with green dash line. G. TaNF-Y complex (co-expression of NF-YA3-D-His, GST-NF-YB7-B and MBP-NF-YC6-B) directly bind to oligo nucleotides containing CCAAT motifs in the *TaGli-γ-700* and *TaLMW-400* promoter via EMSA assay. “+” and “-” indicate the presence and absence of the probe or protein, respectively. EMSA assays using probe derived from the TaNF-Y target gene promoters containing CCAAT motif identified in this study. Unlabeled probe and mutated probe were used as competing DNA fragments. H. The TaNF-Y complex inhibits the expression of *TaGli-γ-700* and *TaLMW-400*, as measured by the dual luciferase reporter assay. Relative LUC/REN value indicates the ratio of the signal detected for firefly luciferase (LUC) versus *Renilla reniformis* luciferase (REN) activity. Data shown mean ± SD of six biological replicates. Statistical significance was determined by Student’s *t* test. **, *P* < 0.01.

Next, we aimed to determine the potential direct targets of TaNF-Y among the DEGs. Leveraging previously generated chromatin accessibility data of the endosperm at 6, 8, and 12 DAP (Zhao et al., 2023), we searched for DEGs with CCAAT motifs present within proximal open chromatin regions. This identified 960 genes (Supplemental Data Set S3), with 514 up-regulated and 446 down-regulated in *TaNF-Y RNAi* lines, including 41 gluten protein coding genes (all up-regulated genes, such as*TaGli-γ-700*, and *TaLMW-400*), and 15 starch biosynthesis genes (six up-regulated genes and nine down-regulated genes) (Figure 3E-F). We further confirmed the directly binding of TaNF-Y to the promoter of certain gluten coding genes, such as *TaGli-γ-700* and *TaLMW-400* via Electronic Mobility Shift Assay (EMSA) (Figure 3G). Notably, TaNF-YA3-D alone did not bind to the promoters of *TaGli-γ-700* and *TaLMW-400*, neither for other components such as TaNF-YB7-B and TaNF-YC6-B (Supplemental Figure S5A), as reported previously in *A. thaliana* (Siriwardana et al., 2016). Whereas, all three components of the TaNF-Y trimeric were required for binding to the promoters of *TaGli-γ-700* and *TaLMW-400* (Figure 3G), suggesting the integrity of the TaNF-Y complex is important. Furthermore, TaNF-YA3-D and TaNF-YC6-B exhibited transcriptional repressive activity in the dual luciferase (LUC) reporter assay in wheat protoplast (Supplemental Figure S5B). LUC reporter assay in tobacco (*N. benthamiana*) leaves further validated the repression of TaNF-Y complex on *TaGli-γ-700* and *TaLMW-400* (Figure 3H). Therefore, TaNF-Y directly represses gluten coding genes to regulate the composition of SSP in wheat.

Although few starch biosynthesis genes are DEG with CCAAT motifs in their open promoter regions (Figure 3E), we did not observe a shift bond in the EMSA assay, such as *TaAGPS1a* promoter (Supplemental Figure S5C). In addition, starch biosynthesis genes generally are down-regulated in *TaNF-YA3* and *TaNF-YC6* attenuation lines (Figure 3C), while TaNF-Y is likely repressor to the targets (Supplemental Figure S5B). These data suggest that TaNF-Y may function separately in regulating the transcription of starch biosynthesis and SSP coding genes.

### TaNF-Y indirectly promotes starch biosynthesis by repressing *TaNAC019*

We speculate that TaNF-Y may facilitate starch biosynthesis indirectly through inhibiting some repressor of starch biosynthesis genes. Among the potential targets of TaNF-Y, *TaNAC019* is reported to negatively regulate starch biosynthesis genetically in wheat (Liu et al., 2020), though other report shows slightly positive regulation of starch biosynthesis gene by TaNAC019 at 24 DAP stage (Gao et al., 2021). *TaNAC019-A/B/D* show lagged activation pattern after decline of *TaNF-YA3-B/D* during endosperm development temporally (Supplemental Figure S6A). Moreover, *TaNAC019-A/B/D* are up-regulated in either *TaNF-YA3-RNAi* or *TaNF-YC6-RNAi* lines at 10 DAP (Figure 4A). In addition, CCAAT motif is present in the proximal open chromatin region of T*aNAC019-B* at 12 DAP (Supplemental Figure S6B). We further confirmed the directly binding of TaNF-Y to the promoter of *TaNAC019-B* via EMSA (Figure 4B), as well as the repression of TaNF-Y to *TaNAC019-B* by LUC reporter assay in tobacco leaf (Figure 4C). Such binding and transcriptional repression is dependent on the CCAAT motif (Figure 4B, C). Thus, *TaNAC019-B* is indeed a direct inhibited target of the TaNF-Y.

**Figure 4.**
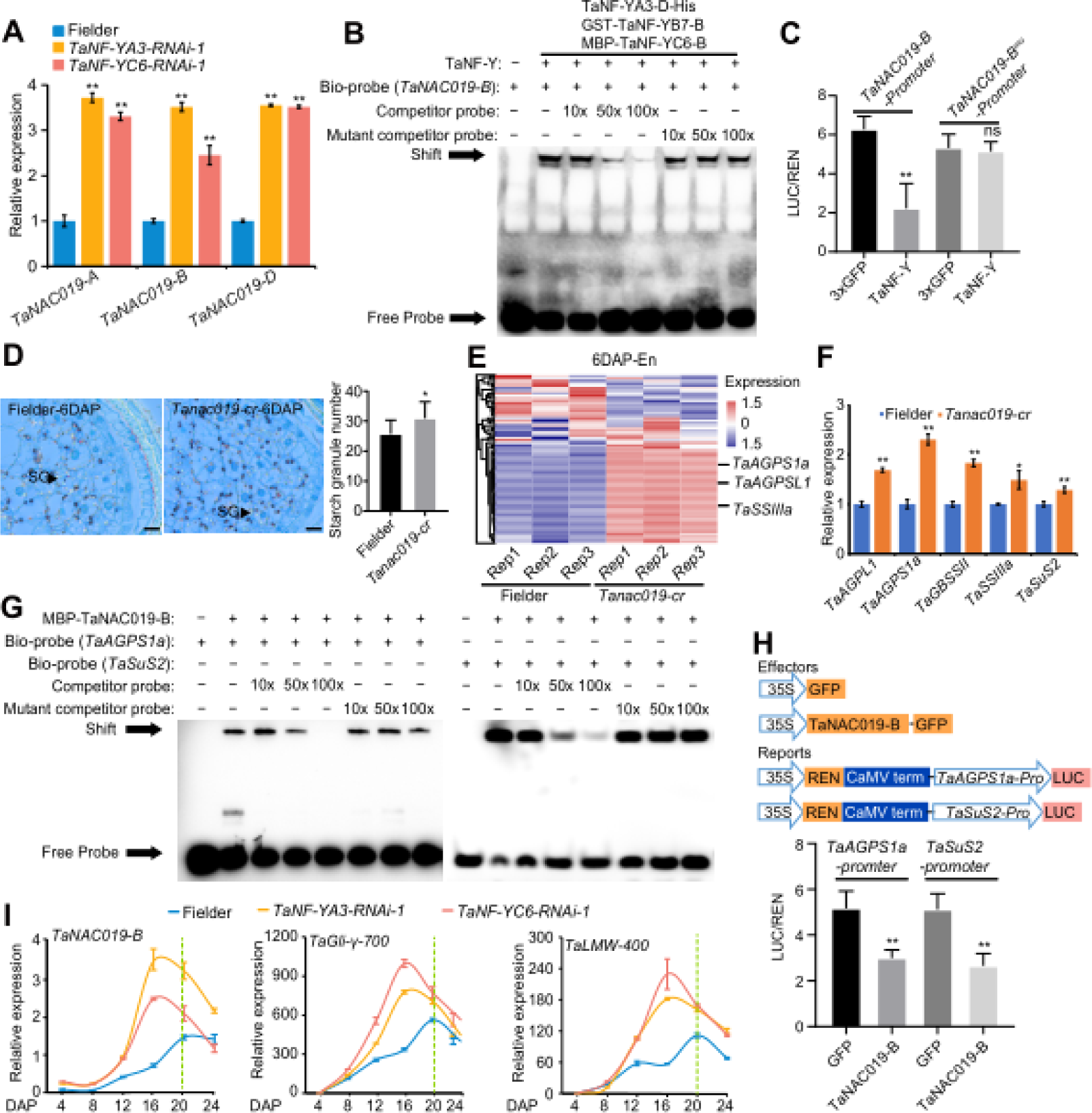
TaNF-Y directly inhibits *TaNAC019* expression and promotes starch biosynthesis. A. Comparison of the expression level of *TaNAC019* triads between Fielder and *TaNF-YA3-RNAi-1* and *TaNF-YC6-RNAi-1* lines. The expression level of genes in Fielder is set as 1, and the relative expression level was normalized to wheat *TaActin*, and reported as mean ±SD of three biological replicates. Statistical significance was determined by Student’s *t* test. **, *P* < 0.01. B. The EMSA assay showing direct binding of TaNF-Y complex (TaNF-YA3-D-His/GST-TaNF-YB7-B/MBP-TaNF-YC6-B) to *TaNAC019-B* promoter sequence. “+” and “-” indicate the presence and absence of the probe or protein, respectively. C. Dual luciferase reporter assay validation of transcriptional inhibition of TaNF-Y complex to *TaNAC019-B*. Mutation of the CCAAT motifs was introduced in the promoter region of *TaNAC019-B*. Statistical significance was determined by Student’s *t* test. **, *P* < 0.01. D. Semi-thin sections of developing Fielder and *Tanac019-cr* seed at 6DAP and the number of starch granules. Scale bar, 100 µm. Statistical significance was determined by Student’s *t* test. *, *P* < 0.05. E. Heatmap showing the expression level of starch biosynthetic genes at 6 DAP developmental endosperm of *Tanac019-cr* and Fielder. F. The relative expression level of *TaAGPL1*, *TaAGPS1a*, *TaGBSSII*, *TaSSIIIa*, and *TaGBSSII* in the *Tanac019-cr*, estimated by qRT-PCR and normalized to *TaActin*. The relative expression level was reported as mean ± SD of three biological replicates. Statistical significance was determined by Student’s *t* test. *, *P* < 0.05; **, *P* < 0.01. G. TaNAC019-B directly bind to the *TaAGPS1a and TaSuS2* promoter via EMSA assay. “+” and “-” indicate the presence and absence of the probe or protein, respectively. Unlabeled probe and mutated probe were used as competing DNA fragments. H. Dual luciferase reporter assay to assess the transcriptional inhibition of TaNAC019-B to *TaAGPS1a and TaSuS2*. Statistical significance was determined by Student’s *t* test. **, *P* < 0.01. I. The expression pattern of the *TaNAC019-B*, *TaGli-γ-700* and *TaLMW-400* during grain development in *TaNF-YA3-RNAi-1*, *TaNF-YC6-RNAi-1* and Fielder. The relative expression level of each gene was normalized to wheat *TaActin* with mean ± SD of three biological replicates. Statistical significance was determined by Student’s *t* test. *, *P* < 0.05; **, *P* < 0.01.

We check the starch contents in *Tanac019-cr* mutant (Gao et al., 2021) in detail as compared to Fielder. Based on semi-thin sections, The *Tanac019-cr* mutant contained more starch compound compared with the Fielder at early endosperm development stage, such as 6 DAP (Figure 4D). We further performed RNA-seq analysis for 6 DAP endosperm of *Tanac019-cr* and Fielder. A total of 1,559 DEGs, including 934 up and 625 down regulated genes has been identified (Supplemental Figure S6C and Supplemental Data Set S4). Notably, a number of starch synthesis genes were up-regulated in *Tanac019-cr* (Figure 4E), suggesting TaNAC019 may repress their expression, as previous report (Liu et al., 2020). This was further validated by qPCR of several starch synthesis genes, such as *TaAGPL1*, *TaAGPS1a*, *TaSuS2* (Figure 4F). These results indicated that TaNF-Y could regulate starch biosynthesis genes’ expression through inhibiting *TaNAC019*.

We further explore how *TaNAC019* affects the transcription of starch synthesis genes. By EMSA assay, we confirmed that TaNAC019-B could bind to the promoter of *TaAGPS1a* and *TaSuS2* (Figure 4G). In addition, we performed transient dual LUC assay in tobacco leaves system. The results showed that TaNAC019-B repress the transcription of *TaAGPS1a* and *TaSuS2* (Figure 4H). Thus, TaNAC019 could directly repress the expression of starch biosynthesis genes, such as *TaAGPS1a* and *TaSuS2,* to negatively regulate starch biosynthesis in particular during early grain filling stage.

Notably, from a time-course qPCR assay, we observed an early shift of peak expression time point of SSP coding genes, such as *TaGli-γ-700* and *TaLMW-400* as well as *TaNAC019-B* in *TaNF-YA3 RNAi-1* and *TaNF-YC6 RNAi-1* lines in addition to elevated expression (Figure 4I). This further indicates that TaNF-Y complex not only regulate the expression level but also the temporal pattern of direct targets during endosperm development.

### TaNF-Y interacts with PRC2 and mediates deposition of H3K27me3 at *TaNAC019* and gluten coding genes

Furthermore, we aimed to investigate how TaNF-Y functions in repressing gene expression. Interestingly, we observed reverse trends between the accumulation of H3K27me3 mark at *TaNAC019-B* and gluten coding genes such as *TaGli-γ-700* and *TaLMW-400* with their expression levels during different endosperm developmental stages (Figure 5A). In *Arabidopsis*, the NF-Y complex was reported to interact with PRC2, the writer complex for H3K27me3, to mediate gene silencing in the context of flowering regulation(Liu et al., 2018b). Several components of the PRC2 complex are highly expressed during endosperm development, including *TaSWN* and *TaFIE* coding genes (Supplemental Figure S7). We investigated the potential protein interaction between PRC2 and the TaNF-Y complex in wheat using Y2H, and BiFC assays (Figure 5B-C). TaSWN-B mainly interacts with TaNF-YC6-B and TaNF-YB7-B in the nucleus (Figure 5B, C).

**Figure 5.**
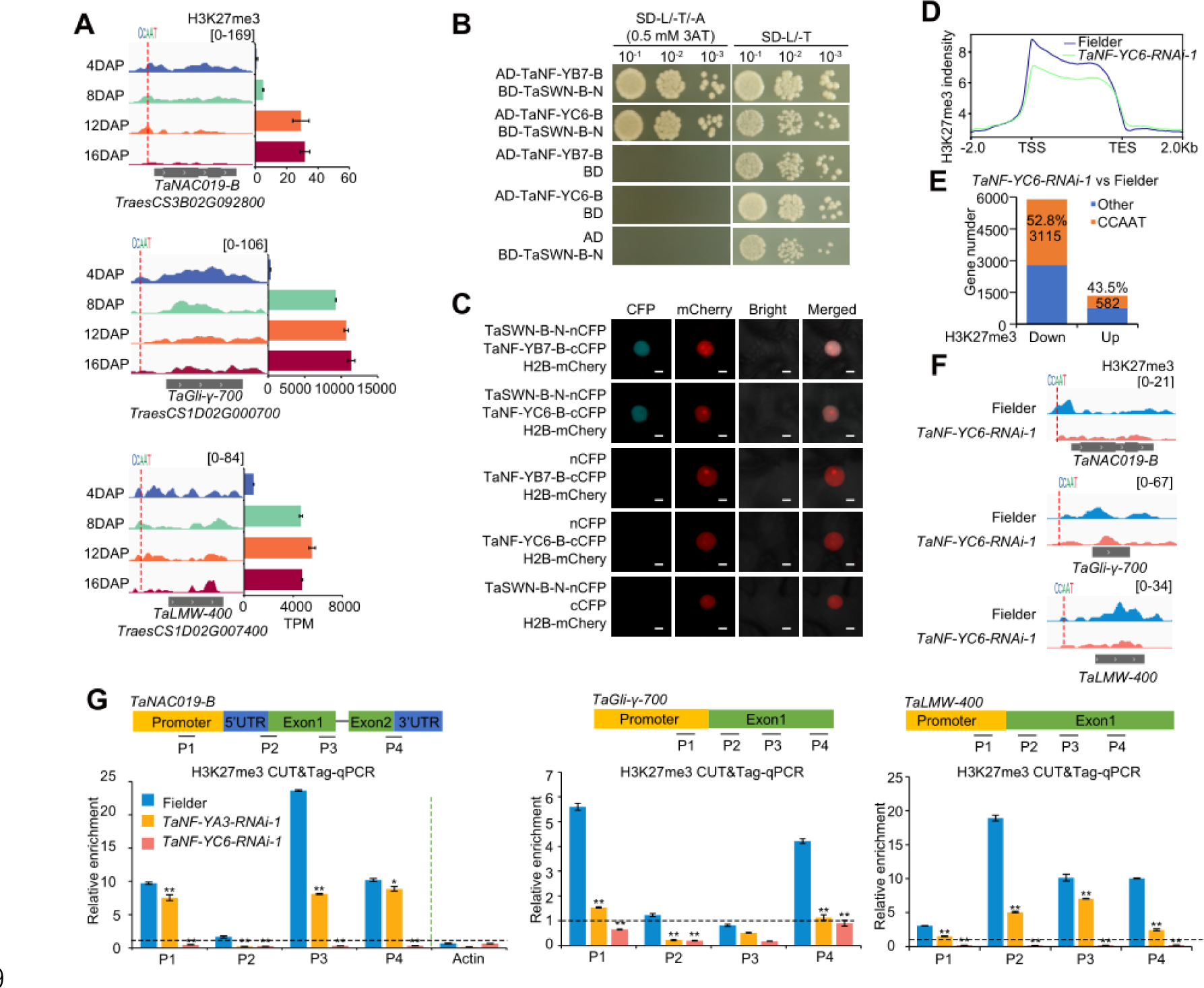
The TaNF-Y interacts with PRC2 to facilitate H3K27me3 deposition for inhibiting *TaNAC019-B* and SSP coding genes. A. IGV shows the dynamic change of H3K27me3 and the transcription level of *TaNAC019-B*, *TaGli-γ-700* and *TaLMW-400* during endosperm development. The CCAAT motif was indicated with red dash line. B. Y2H assay showing the interaction of TaNF-YB7-B and TaSWN-B-N, TaNF-YC6-B and TaSWN-B-N. C. BiFC assay showing the interaction between TaNF-YB7-B and TaSWN-B-N, TaNF-YC6-B and TaSWN-B-N in tobacco leaves. H2B-mCherry was used as transformation control and indication of nuclei. Scale bar, 0.5 µm. D. The meta profile of H3K27me3 along genes in *TaNF-YC6-RNAi-1* and Fielder. E. The number of genes with altered H3K27me3 modification and open containing CCAAT motif in the *TaNF-YC6-RNAi-1* line compared to Fielder. F The IGV showing the H3K27me3 level of *TaNAC019-B*, *TaGli-γ-700*, and *TaLMW-400* loci in Fielder and *TaNF-YC6-RNAi-1*. The CCAAT motif was indicated with red dash line. G. CUT&Tag-qPCR verification the H3K27me3 modification difference at *TaNAC019-B*, *TaGli-γ-700*, and *TaLMW-400* loci between *TaNF-YA3-RNAi-1, TaNF-YC6-RNAi-1* and Fielder. The H3K27me3 fold change value was normalized to input. A value > 1 indicates enrichment of binding sites in that region whereas a value < 1 indicates relative depletion of binding sites *TaActin* was used as a negative control. Data showing mean ± SD (n = 3). Statistical significance was determined by Student’s *t* test. *, *P* < 0.05; **, *P* < 0.01.

Whether such interaction influence the deposition of H3K27me3 at TaNF-Y targets? To answer this question, we conducted Cleavage Under Targets & Tagmentation (Cut&Tag) (Zhao et al., 2023) to obtain whole genome-wide H3K27me3 profiles in *TaNF-YC6-RNAi-1* line and Fielder, since TaNF-YC6-B interacts with TaSWN-B. There was a general reduction of H3K27me3 peak coverage and numbers in *TaNF-YC6-RNAi-1* line compared to Fielder (Supplemental Figure S8). The mega profile of H3K27me3 at gene body regions is declined in *TaNF-YC6-RNAi-1* line compared to Fielder (Figure 5D). Genes containing CCAAT motifs showed a dramatic reduced H3K27me3 levels than increased in *TaNF-YC6-RNAi-1* line (Figure 5E and Supplemental Data Set S5), including *TaNAC019-B*, *TaGli-γ-700*, and *TaLMW-400* (Figure 5F).

We further performed CUT&Tag-qPCR to quantitatively measure the accumulation of H3K27me3 at *TaNAC019-B*, *TaGli-γ-700*, and *TaLMW-400* loci using endosperm tissue from both *TaNF-YA3* and *TaNF-YC6 RNAi* lines as compared to Fielder. A significant reduction of H3K27me3 level at regulatory regions and gene body of both genes was observed in *TaNF-YA3-RNAi-1* and *TaNF-YC6-RNAi-1* lines compared to Fielder (Figure 5G). These findings suggest that the TaNF-Y complex interacts with PRC2 components and mediates the deposition of H3K27me3 at *TaNAC019-B* and gliadin coding genes to properly regulate their temporal expression patterns during endosperm development.

### Mutation of PRC2 influences starch synthesis and gluten protein composition

Considering the significance of H3K27me3 in silencing the expression of *TaNAC019-B* and gliadin coding genes, we conducted further investigations on the endosperm developmental defects of PRC2 core component mutant. Since PRC2-mediated deposition of H3K27me3 is crucial for wheat embryo development (Zhao et al., 2023). we focused on the genome-edited *Tafie-C87* line, which exhibited slightly delayed embryo development but produced fertile seeds. This line carried frameshift mutations in the A and D subgenomes of *TaFIE* (*TraesCS7A02G308300*, *TraesCS7D02G305100*) and a two-amino acids deletion in the B subgenome (*TraesCS7B02G377900LC*) (Zhao et al., 2023).

The *Tafie-C87* line displayed altered seed size with decreased GW, increased GL, and a significant reduction in TGW compared to the control KN199 (Figure 6A, B). Consistent with the shriveled seed, the *Tafie-C87* line exhibited a significant decrease in total starch content (Figure 6C) and SG number, as indicated by coomassie brilliant blue staining of semi-thin sections (Figure 6D and Supplemental Figure S9A). SEM analysis also revealed the diameter of A-Type SGs and the number of B-Type SGs are both decreased in the *Tafie-C87* mature seeds (Supplemental Figure S9B, C). Furthermore, the amount of total protein increased in *Tafie-C87* (Figure 6E). In addition, only gliadins contents were significantly increased in *Tafie-C87*, the level of HMW-GS and LMW-GS were similar to KN199 (Figure 6F, G).

**Figure 6.**
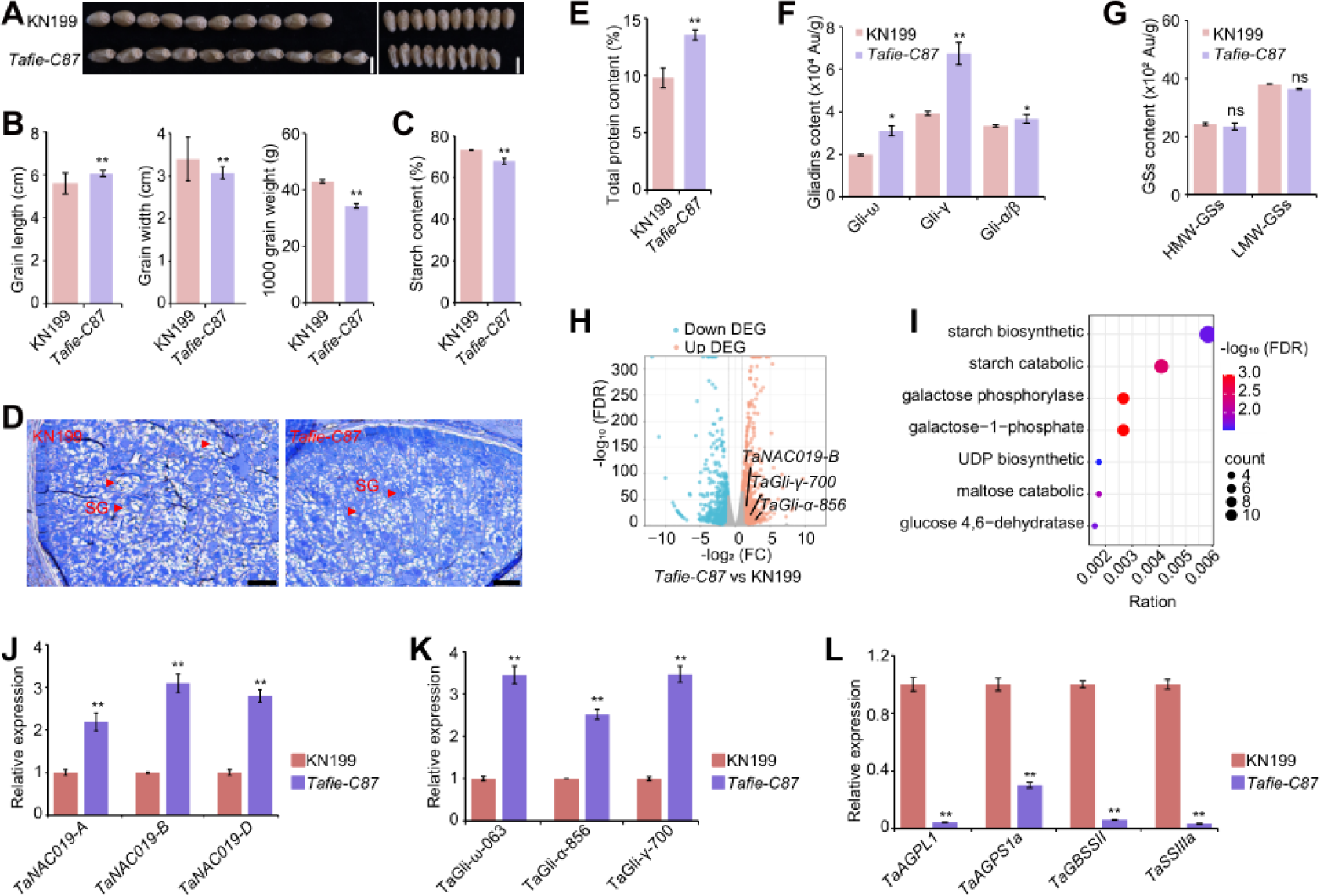
Mutation of TaFIE influences starch synthesis and SSP composition. A-G. Grain morphology (A), and the statistical comparisons of grain size-related traits (B), starch content (C). histological section analysis of KN199 and *Tafie-C87* endosperm starch (D). Total protein content (E), gliadins content (F) and GSs content (G) between KN199 and *Tafie-C87*. Three grains were used for histological analysis and starch granules staining with coomassie brilliant blue. Starch granules (SG) were indicated with red triangles, scale bar, 100 µm. The Grain size-related traits were determined using a camera-assisted phenotyping system with five biological replicates, each with approximately 10 g seeds. The contents of HMW-GS, LMW-GS, and gliadin were estimated by RP-HPLC, analysis with 3 biological replicates. Statistical significance was determined by Student’s *t* test. *, *P* < 0.05; **, *P* < 0.01, ns, no significant difference. H. Volcano plot showing the up- and down-regulated genes of the *Tafie-C87* compare with KN199. I. GO enrichment analysis of down-regulated genes in the *Tafie-C87*. J-L. The relative expression level of *TaNAC019* (J), gliadins coding genes (K), and starch synthesis coding genes (L) in *Tafie-C87* and KN199. The relative expression level of each gene was estimated by RT-qPCR using 10 DAP endosperm, normalized to wheat *TaActin*, and reported as mean ± SD of three biological replicates. Statistical significance was determined by Student’s *t* test. *, *P* < 0.05; **, *P* < 0.01.

Consistent with the phenotypical defects, RNA-seq analysis of developing endosperm of KN199 and *Tafie-C87* line further revealed that numerous genes expression altered, including up-regulation of *TaNAC019-B*, *TaGli-γ-700*, *TaGli-α-856* (Figure 6H, Supplemental Data Set S6). Notably, the *Tafie-C87* line displayed a significant reduction of genes encoding enzymes involved in starch biosynthesis and catabolic process (Figure 6I). The up-regulation of *TaNAC019-A/B/D*, and SSP related genes *TaGli-ω-063*, *TaGli-α-856* and *Tali-γ-700*, as well as down-regulation of starch biosynthesis genes *TaAGPL1*, *TaAGPS1a*, *TaGBSSII*, *TaSSIIIa* were confirmed by qPCR assay in *Tafie-C87* line compared to KN199 (Figure 6J-L). These findings confirm the involvement of PRC2 in the regulation of starch synthesis and gluten protein composition through transcriptional regulation.

### Natural variation of TaNF-Y component is associated with starch and protein contents for breeding selection

Grain size and end-use quality are crucial traits that have been subject to domestication and breeding selection (Michel et al., 2019). To evaluate the contribution of the TaNF-Y components in breeding selection, we conducted natural variation analysis in the genic region of *TaNF-YA*, *TaNF-YB*, and *TaNF-YC*, utilizing the Watkins datasets (Cheng et al., 2023). In general, we found variation of four TaNF-YAs, eight TaNF-YBs, four TaNF-YCs, were associated with seed developmental traits (Supplemental Data Set S7). Among them, ten genes are linked to TGW, four genes are linked to grain protein contents, and six genes are linked to grain starch contents (Supplemental Data Set S7). In particular, we identified six SNPs in the coding region and downstream of *TaNF-YB7-B* (*TraesCS7B02G121900*), which defined two haplotypes (Hap1 and Hap2) with the frequency of 43.4% and 56.6%, respectively (Figure 7A). Significant differences in TGW, grain protein and starch content values were observed between Hap1 and Hap2 (Figure 7B), with Hap2 exhibiting higher grain starch contents but lower protein contents. The frequency of Hap2 in modern cultivars (92.5%) was significantly higher than that in landraces (47.9%) (Figure 7C), indicating Hap2 was selected during modern wheat breeding.

**Figure 7.**
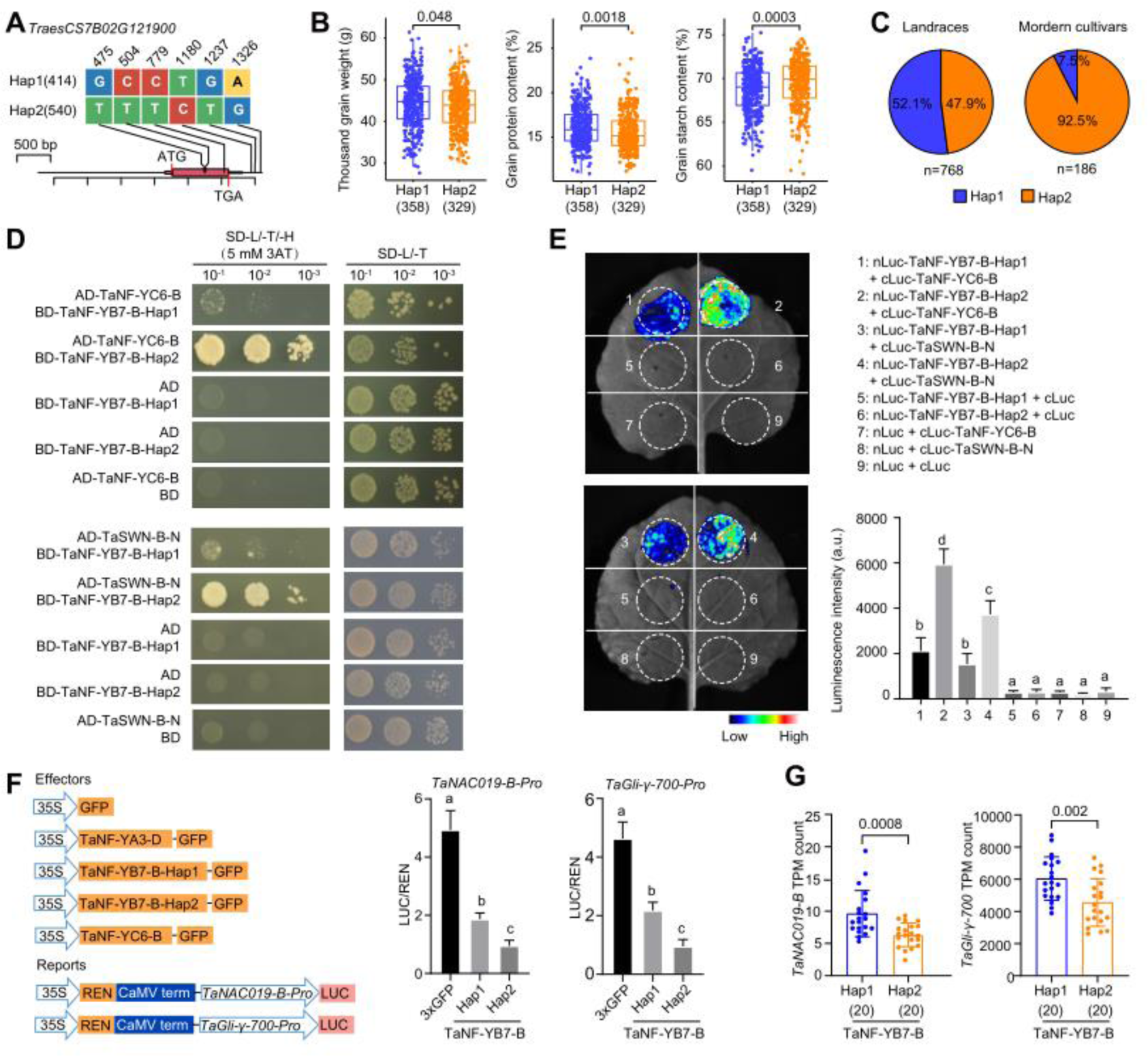
Natural variation of *TaNF-YB7-B* affects targets inhibition and is associated with starch and protein contents for breeding selection. A. Schematic diagram showing the variations in the upstream, coding regions, and downstream region of *TaNF-YB7-B* in the wheat population including 768 Watkins landraces and 186 modern cultivars. B. Comparison of thousand-grain weight, grain protein content, and starch content between TaNF-YB7-B Hap1 and Hap2. Statistical significance was determined using the Wilcoxon rank-sum test. The numbers in parentheses represent the sample size used for statistical analysis. Statistical significance was determined by Student’s *t* test. C. Distribution of different haplotypes of TaNF-YB7-B in landraces and modern cultivars. *n* indicates the sample size. D. Y2H shows the interaction between TaNF-YC6-B, TaSWN-B-N and TaNF-YB7-B-Hap1, TaNF-YB7-B-Hap2, respectively. E. SPLC assay. nLUC-tagged TaNF-YB7-B-Hap1 or nLUC-tagged TaNF-YB7-B-Hap2 was co-transformed into tobacco leaves along with cLUC-tagged TaNF-YC6-B or TaSWN-B-N, the numbers in the circles represent different plasmid combinations. Values are means ± SD (n = 3). The different letters above each bar indicate that means differ significantly by one-way ANOVA and Tukey’s multiple comparison test at *P* < 0.05. F. Dual luciferase reporter assay to assess the transcriptional inhibition of different haplotypes of *TaNF-YB7-B* in regulation of *TaNAC019-B*, and *TaGli-γ-700*. The different letters above each bar indicate that means differ significantly by one-way ANOVA and Tukey’s multiple comparison test at *P* < 0.05. G. *TaNAC019-B*, and *TaGli-γ-700* expression with wheats groupd by the *TaNF-YB7-B* two different haplotypes. Statistical significance was determined by Student’s *t* test (n = 20 for Hap1; n = 20 for Hap2).

To understand how the DNA variations within *TaNF-YB7-B* coding region contribute to the functional diversity of the protein product, we conducted interaction tests among TaNF-Y trimeric components and its interaction with PRC2 using different coding sequences of *TaNF-YB7-B* (Figure 7D). Y2H revealed TaNF-YB7-B-Hap2 exhibiting higher interaction intensity with TaNF-YC6-B, as well it interacts with TaSWN-B stronger than TaNF-YB7-B Hap1 (Figure 7D). The higher interaction intensity between TaNF-YB7-B-Hap2 and TaNF-YC6-B, TaSWN-B was further confirmed by split luciferase complementation (split-LUC) assay (Figure 7E). Since all three components of TaNF-Y are required for the binding of TaNF-Y to targets (Figure 4) and TaNF-Y interacts with PRC2 to repress gene expression (Figure 5), we further evaluated the effect of different coding haplotypes of *TaNF-YB7-B* on the transcriptional regulation of TaNF-Y to targets using a LUC reporter assay (Figure 7F). The results demonstrated that the coding variations in *TaNF-YB7-B* indeed affect the repressive transcriptional activity. The Hap2 variant exhibited a higher repression effect on TaNF-Y targets, such as *TaNAC019-B*, and *TaGli-γ-700* (Figure 7F). Consistently, *TaNAC019-B* and *TaGli-γ-700* exhibited significantly higher expression levels in the developing grain (20 DAP) of accessions with TaNF-YB7-B-Hap1 than that of Hap2 (Figure 7G), according to the population-wide transcriptome (Zhao et al., 2023). This further validates that *TaNF-YB7-B-Hap2* increases starch and decreases protein content in endosperm associated with repressing the expression of *TaNAC019-B* and *TaGli-γ-700*.

Thus, natural variations in TaNF-Y components can influence starch biosynthesis and gluten protein composition by modulating transcriptional regulation. These variations have been selected during the modern breeding process.

## Discussion

Balancing high grain yield and superior seed quality is a persistent challenge in crop improvement endeavors (Michel et al., 2019). This challenge likely arises from the intricate (Pleijel and Uddling, 2011). Achieving this balance necessitates a deep understanding of the molecular mechanisms governing starch and SSP biosynthesis and the identification of regulatory factors that can be manipulated to separately optimize both aspects.

### Temporal expression of starch and SSP biosynthesis genes: epigenetic regulation and key TFs

The proper development of starch biosynthesis and gluten protein accumulation in the endosperm is notably reflected in the distinct temporal expression patterns of related genes (Figure 1). However, the mechanisms underlying the generation of these expression patterns remain largely enigmatic, although several upstream TFs have been identified. A notable observation is the presence of a high level of repressive histone modification, H3K27me3, surrounding SSP-encoding genes before their activation at 6 DAP. Subsequently, as SSP activation occurs during later endosperm development (8-16 DAP), H3K27me3 levels decline (Figure 5). While the causal relationship between H3K27me3 and transcriptional regulation remains elusive, we did observe the activation of key SSP genes in a partially *TaFIE* (PRC2 component) -knockout mutant line (Figure 6). Notably, the *Tafie-C87* mutant line exhibited elevated gliadins and total protein contents. Conversely, starch biosynthesis-related genes displayed minimal H3K27me3 coverage throughout endosperm development, indicating a lesser influence of histone modification. Instead, TaNAC019, a known repressor of starch biosynthesis (Liu et al., 2020), and further validated its direct repression to starch biosynthetic genes here (Figure 5), exhibited an expression pattern opposite to that of starch biosynthetic genes. Interestingly, *TaNAC019-B* appeared to be a suitable target for H3K27me3, as H3K27me3 levels declined after 8 DAP while *TaNAC019-B* expression gradually increased (Figure 5). Similar to SSP genes, *TaNAC019-B* was up-regulated in the *Tafie-C87* mutant line (Figure 6).

These findings suggested an association between epigenetic modifications, particularly H3K27me3, and the temporal expression patterns of SSP and starch-related genes, although the precise influence may be direct or indirect. This discovery expands our understanding of the role of PRC2-H3K27me3 in regulating later endosperm development, in addition to its known involvement in the initiation of endosperm proliferation and cellularization (Xiao et al., 2015, Zhang et al., 2023)

### The multi-face role of the NF-Y complex in starch and SSP biosynthesis

The NF-Y complex has been recognized for its involvement in seed development across different plant species primarily through its transcription factor activity (Laloum et al., 2013). Among the three subunits of the NF-Y complex, NF-YA typically mediates binding to the CCAAT motif, while NF-YB or NF-YC may interact with other TFs or chromatin modification factors (Myers and Holt, 2018). In our study, we identified an endosperm core TaNF-Y complex that functions as a repressor by interacting with PRC2 and facilitating the deposition of H3K27me3 at target loci (Figure 5). TaNF-Y directly binds to SSP-encoding genes, ensuring specific enrichment of H3K27me3 by recruiting PRC2 to maintain a repressed transcriptional state of SSP genes during early endosperm development. This repressive chromatin status diminishes as TaNF-Y complex levels decline with endosperm development progression, aligning with the activation of SSP genes at later stages. Notably, in *TaNF-YA3-RNAi-1* and *TaNF-YC6-RNAi-1* lines, not only did the expression levels of *TaGli-γ-700* and *TaLMW-400* increase, but their peak expression times also advanced by several days (Figure 4).

Intriguingly, TaNF-Y did not bind to some starch biosynthesis enzyme-encoding genes, even when CCAAT motifs were present in their promoters (Supplemental Figure S4). Instead, TaNF-Y directly inhibited the expression of *TaNAC019-B*, which displayed a “perfect” lagged activation upon the decline of *TaNF-Y* expression (Figure 1). Furthermore, *TaNAC019-B* exhibited elevated expression with an earlier peak when *TaNF-Y* was knocked down (Figure 4). By integrating both direct and indirect pathways, TaNF-Y is intricately involved in the regulation of both starch and SSP in wheat.

### Decoupling starch and SSP biosynthesis for yield and quality improvements

Natural variations within the components of the TaNF-Y complex present promising opportunities for discovering elite alleles. Indeed, we identified two amino acid variations within the coding region of *TaNF-YB7-B*, which influence the formation of the TaNF-Y trimeric complex and its interaction with TaSWN. Consequently, different haplotypes of TaNF-Y exhibit varied transcriptional repressive activities towards downstream targets, such as *TaNAC019* and SSP genes. This variability is linked to altered starch and protein contents, ultimately influencing grain yield and seed quality (Figure 7). The distinct regulation pathways for starch and SSP by the TaNF-Y-PRC2 module provide a unique avenue for decoupling these two processes. Further analyses may uncover additional variations of TaNF-Y that specifically influence either starch or SSP biosynthesis. With the advances in molecular breeding techniques, including genome editing, these findings open promising avenues for the development of crop varieties that simultaneously enhance grain yield and seed quality.

In summary, our study identifies a core TaNF-Y complex that plays a pivotal role in regulating both starch and gluten protein contents in wheat. This regulation is associated with specific PRC2-mediated deposition of H3K27me3 modifications on key factors associated with SSP biosynthesis and the critical TF, TaNAC019, which regulates key enzymes involved in starch biosynthesis. Importantly, the natural variation of the *TaNF-Y* component *TaNF-YB7-B* is identified to associate with starch and protein contents in a panel of elite cultivars in China and selected during breeding process (Supplemental Figure S10).

## Methods

### Plant materials and growth conditions

For *TaNF-YA3-RNAi* and *TaNF-YC6-RNAi* transgenic lines (in the Fielder background), the conserved fragments of *TaNF-YA3* (380bp), *TaNF-YC6* (300bp) were cloned into PC414C and inserted into the RNAi vector PC336 as inverted repeats to create the *TaNF-YA3-RNAi* and *TaNF-YC6-RNAi* recombinant vectors using Gateway technology (Wang et al., 2022). The recombinant constructs were transformed into *Agrobacterium tumefaciens* strain EHA105. The transgenic wheat lines were generated through Agrobacterium-mediated infiltration of immature wheat embryos as previously described (Ishida et al., 2015). The resulting T0 to T2 transgenic lines were selected via PCR using ubiquitin promoter-specific primers (Supplemental Data Set S8). For the *TaNAC019* mutants, we used the *Tanac019-cr* line as previously described (Gao et al., 2021). For the *Tafie* mutants, we use the *Tafie-C87* line as previously described (Zhao et al., 2023). All wheat plants were grown in the experimental field of Beijing (39 55′ N, 116 23′ E) and in a greenhouse under long-day conditions (16 h light–8 h dark cycles). *N*. *benthamiana* was grown in a greenhouse at 22 °C under a 16 h light and 8 h darkness photoperiod.

### Yeast two-hybrid assay

Yeast two-hybrid assays were performed as described in the Frozen-EZ Yeast Transformation II™ (Zymo Research). For the interaction of TaNF-YA3-D, TaNF-YB7-B and TaNF-YC6-B, the coding region sequence of *TaNF-YA3-D*, *TaNF-YB7-B* were introduced into the prey vector (pGADT7). To construct the bait vector, we ligated the full-length CDSs of *TaNF-YC6-B* into the bait vector (pGBKT7) vector. For the interaction of TaNF-YB7-B, TaNF-YC6-B with TaSWN-N, we ligated the full-length CDSs of *TaNF-YB7-B*, *TaNF-YC6-B* into the prey vector (pGADT7) vector and *TaNWN-B-N* into the bait vector (pGBKT7). For the interaction of TaNF-YC6-B, TaSWN-B-N with TaNF-YB7-B-Hap1 and TaNF-YB7-B-Hap2, respectively, we cloned the CDS of *TaNF-YC6-B* and *TaSWN-B-N* into the prey vector (pGADT7), *TaNF-YB7-B-Hap1* and *TaNF-YB7-B-Hap2* into the bait vector (pGBKT7), and selected on DDO (Synthetic Dropout Medium/-Tryptophan-Leucine) and TDO (Synthetic Dropout Medium/-Tryptophan-Histone-Leucine) media (Clontech) with corresponding concentration of 3-AT. The empty pGADT7 or empty pGBKT7 used as a negative control. The primers are listed in Supplemental Data Set S8.

### Bimolecular fluorescence complementation (BiFC) assay

For the construction of BiFC vectors, pSCY-NE(R)-nCFP is the vector for N-terminal fusion to cyan fluorescent protein (nCFP), and pSCY-CE(R)-cCFP is the vector for C-terminal fusion to CFP (cCFP). For the interaction of TaNF-YA3-D, TaNF-YB7-B and TaNF-YC6-B, the full-length CDS of *TaNF-YA3-D*, and *TaNF-YB7-B* were cloned into pSCY-NE(R)-nCFP. The full-length CDS of *TaNF-YC6-B* was cloned into pSCY-CE(R)-cCFP. For the interaction of TaSWN-B-N with TaNF-YB7-B and TaNF-YC6-B, the N-terminal of *TaNWN-B* (1050 bp) was cloned into pSCY-NE(R)-nCFP. The CDS of *TaNF-YB7-B* and *TaNF-YC6-B* were cloned into pSCY-CE(R)-cCFP. Young leaves of 4-week-old *N. benthamiana* plants were co-infiltrated with Agrobacterium (strain GV3101) harbouring different combinations of these plasmids. *N. benthamiana* plants grown in long-day conditions for 48 h after infiltration, the CFP signals were detected by a confocal microscope (LSM980; Carl Zeiss, Germany). H2B-mCherry was used as a cell nucleus marker, The primers are listed in Supplemental Data Set S8.

### Phylogenetic analysis

We obtained NF-YA, NF-YB, and NF-YC subunit protein sequences of *A. thaliana*, *O. sativa* and *T. aestivum* from Plant Transcription Factor Database (PlantTFDB) (version 5.0, http://planttfdb.gao-lab.org/), We aligned the amino acid sequences with ClustalW and constructed a phylogenetic tree using the neighbour-joining method in MEGA 11 software with default parameters, The evolutionary distances were calculated using the Poisson model. The phylogeny test was computed using bootstrap method with 10,000 replications.

### Subcellular localization

The full-length open-reading frames (ORFs) of *TaNF-YA3-D*, *TaNF-YB7-B* and *TaNF-YC6-B* were amplified using specific primers and cloned in the pBI221-EGFP vector containing the *CaMV 35s* Promoter. The resulting construct *p35s*-*TaNF-YA3-D*-EGFP, *p35s TaNF-YB7-B*-EGFP, *p35s TaNF-YC6-B*-EGFP and the *p35s* EGFP served as a control were separately introduced into wheat protoplasts. The transformed protoplasts were cultured at 22°C under darkness for 18 h, and GFP signals observed on a confocal microscope (LSM980; Carl Zeiss, Germany). The primers are listed in Supplemental Data Set S8.

### Construction of transgenic wheat lines

For the RNAi construct, conserved fragments of *TaNF-YA3* (380bp), *TaNF-YC6* (300bp) were cloned into PC414C and inserted into the RNAi vector PC336 as inverted repeats to create the *TaNF-YA3-RNAi* and *TaNF-YC6-RNAi* recombinant vectors using Gateway technology (Wang et al., 2022). The recombinant constructs were transformed into *Agrobacterium tumefaciens* strain EHA105. The transgenic wheat lines were generated through Agrobacterium-mediated infiltration of immature wheat embryos as previously described (Ishida et al., 2015). The resulting T0 to T2 transgenic lines were selected via PCR using ubiquitin promoter-specific primers (Supplemental Data Set S8). For the *TaNAC019* mutants, we used the *Tanac019-cr* line as previously described (Gao et al., 2021). For the *Tafie* mutants, we use the *Tafie-C87* line as previously described (Zhao et al., 2023).

### Phenotypic analysis

The seed-related phenotypes including TGW, grain length and width were determined using a camera-assisted phenotyping system (Wanshen Detection Technology Co., Ltd, Hangzhou, China). To determine total starch content, 100 mg flour was used and measured using Megazyme (Irishtown, Ireland) Total Starch Assay Kit (catalog: K-TSTA) according to the manufacturer’s instructions (Botticella et al., 2018). The contents of HMW-GSs, LMW-GSs, and gliadins were detected by RP-HPLC as previously described (Gao et al., 2021). Total protein content was evaluated using the voluntary standard method of the National Standard of China 5506.4-2008 (GB/T5506.4-2008).

### Semi-thin sections and scanning electron microscopy

For Fielder, *TaNF-YA3-RNAi*, *TaNF-YC6-RNAi*, KN199 and *Tafie*-*C87*, seeds at 10 DAP were sampled from spikes. For *Tanac019-cr* mutant, seeds at 6 DAP were selected. Seeds were fixed in FAA solution (63% ethanol, 5% acetic acid, 2% formaldehyde) under vacuum for 2 hours and stored at 4 °C for use. The samples were dehydrated through a graded ethanol series and embedded in Technovit 7100 resin (Kulzer, https://www.kulzer-technik.de), according to the manufacturer’s instructions. Sections (2-µm thick) were cut using an UC7&2265 microtome (Leica). To visualize starch granules and protein bodies, the sections were stained with 0.1% w/v coomassie brilliant blue R-250 (Tonosaki et al., 2021). Images of stained sections were captured using an OLYMPUS DP74 Microscope. Endosperm starch granules number was manually calculated for three seeds and the differences between Fielder *TaNF-YA3-RNAi-1*, *TaNF-YC6-RNAi-1*, *Tanac019-cr*, KN199 and *Tafie-C87* were by one-way ANOVA.

For scanning electron microscopy observation, the grains were dried at 37°C for 1 week after harvest. The seeds of *TaNF-YA3-RNAi*, *TaNF-YC6-RNAi*, Fielder, *Tafie-C87* and KN199 were broken transversely, and the ruptured surfaces were coated with gold. Crossbeam 340 & VCT500 (Carl Zeiss, Germany) scanning electron microscope was employed to observe the samples.

### RNA extraction, qRT-PCR analysis, and RNA-seq

Total RNAs were extracted from endosperm at 4 DAP, 8 DAP, 10 DAP, 12 DAP, 16 DAP, 20 DAP, 24 DAP of Fielder, *TaNF-YA3-RNAi-1*, *TaNF-YC6-RNAi-1*, 10 DAP of KN199 and *Tafie-C87*, 6 DAP of Fielder and *Tanac019-cr* using the Quick RNA Isolation Kit with on-column DNaseI digestion (Huayueyang, China) according to the manufacturer’s protocol. First-strand cDNAs were generated using a reverse transcription kit (TRANSGEN, KR116-02). Subsequent qRT-PCR assays were performed using the SYBR Green PCR Master Mix (Vazyme Biotech, Q121-02/03). with each qRT-PCR assay being replicated at least three biological replicates and three technical replicates. Wheat *Actin* gene (*TaActin*, *TraesCS5A02G124300*) was used as an internal reference. Relevant primer sequences are given in Supplemental Data Set S9. For RNA sequencing, oligo (dT) was used for enriching the mRNA from total RNA and then fragmentation and random primer was used for reverse transcript process. Sequencing is performed via the Illumina NovaSeq platform by Annoroad Gene Technology (Li et al., 2022).Clean reads were aligned to IWGSC RefSeq v1.1 (https://wheat-urgi.versailles.inrae.fr/Seq Repository/Assemblies) using hisat2 (v2.0.5) (International Wheat Genome Sequencing, 2018; Kim et al., 2019) The expected number of transcripts per kilobase million (TPM) was used for estimating gene expression levels. Absolute value of Log2 Fold Change ≥ 1 and FDR ≤ 0.05 were considered to be differentially expressed genes.

### Transcriptional activity analysis

To test the transcription activity, the coding sequence of *TaNF-YA3-D* and *TaNF-YC6-D* were cloned into the 35s-GAL4-BD to generate effectors. Virion protein 16 (VP16) contain an acidic transcriptional activation domain was used as a control to identify the effect of TaNF-YA3-D and TaNF-YC6-B on transcriptional activity driven by the UAS module in the reporter vector 35sLUC. The reference vector pRTL containing a *35S* promoter-driven REN was used as an internal control. Each effector construct together with the reporter and reference vectors were co-transformed into wheat protoplasts (Bart et al., 2006; Yoo et al., 2007). The GLOMA 20/20 LUMINOMETER detector (Promega) was used to measure fluorescence signals. Relative LUC activity was calculated by the ratio of LUC/REN. Each sample was tested in four replications. The primers are listed in Supplemental Data Set S8.

### Dual-luciferase reporter assay

To generate *pTaNAC019-B*: LUC, *pTaGli-γ-700*-LUC, *pTaLMW-400*: LUC, *pTaAGPS1a*: LUC and *pTaSuS2*: LUC constructs, we amplified 2-Kb promoter fragments upstream of each gene from CS and ligated them with the CP461-LUC as the reporter vector. The *p35s*-TaNF-YA3-D-GFP, *p35s*-TaNF-YB7-B-GFP, *p35s*-TaNF-YC6-D-GFP and *p35s*-TaNAC019-B-GFP constructs were used as effectors and these plasmids were transformed into GV3101. Then these strains were injected into *N. benthamiana* leaves in different combinations. Dual luciferase assay reagents (Promega, VPE1910) with the Renilla luciferase gene as an internal control were used for luciferase imaging. The Dual-Luciferase Reporter Assay System kit (Cat#E2940, Promega) was used to quantify fluorescence signals. Relative LUC activity was calculated by the ratio of LUC/REN. The relevant primers are listed in Supplemental Data Set S8.

### Protein expression and purification

We cloned the *TaNF-YA3-D* coding sequences into the pET-32a vector (Novagen), *TaNF-YB7-B* coding sequences into the pGEX-4T-1 (Amersham), *TaNF-YC6-B* and *TaNAC019-B* coding sequence into the pMAL-c5X-MBP (Ashutosh Chilkoti). The recombinant proteins were expressed in Escherichia coli BL21 (DE3). Induction was performed by adding IPTG to a final concentration of 0.5 mM and cells were cultured at 28°C for an additional 16 h. Recombinant proteins were purified using the His-tag Protein Purification Kit (Ni-NTA Agarose, QIAGEN GmbH), GST-tag Protein Purification Kit (Glutathione Sepharose, BioVision) and MBP-tag Protein Purification Kit (Dextrin Beads, Smart-Lifesciences), respectively, and were later used for pull-down and EMSA. The primers are listed in Supplemental Data Set S8.

### Pull-down assay and Immunoblot analysis

The recombinant GST-TaNF-YB7-B protein was purified and immobilized on Glutathione Sepharose beads following the manufacturer’s instructions. The beads were divided into two equal aliquots and incubated with the same amount of TaNF-YA3-D-His protein lysate, together with MBP-TaNF-YC6-B or with MBP for 2 hours at 4°C. The beads were subsequently washed five times with PBST buffer, followed by elution with 100 μl of elution buffer (50 mM Tris-HCl, 10 mM reduced glutathione, pH 8.0). Supernatants were resolved by 12% SDS-PAGE and subjected to immunoblotting using anti-GST (LABLEAD, G1001, 1:2500), anti-His (EASYBIO, BE2019-100, 1:2500), anti-MBP (Proteintech, 66003-1-Ig, 1:2500).

### Electrophoretic mobility shift assay (EMSA)

EMSA analyses were performed essentially as previously described (Siriwardana et al., 2016). The double-stranded probes (labeled with biotin at their 5’-end) were annealed from complementary oligonucleotides by cooling from 100°C to room temperature in annealing buffer (10mM Tris, pH 7.5 - 8.0, 50mM NaCl, 1mM EDTA). The sequences of the probes are listed in Supplementary table 7. DNA binding reactions (20μl) (20nM probe, 12mM Tris-HCl pH 8.0, 50mM KCl, 62.5mM NaCl, 0.5mM EDTA, 5mM MgCl_2_, 2.5mM DTT, 0.2 mg/ml BSA, 5% glycerol, 6.25ng/μl poly dA-dT) were incubated TaNF-YA3-D-His/GST-TaNF-YB7-B/MBP-TaNF-YC6-B trimers, three recombinant proteins were pre-mixed in room temperature for 1 hour, then added to DNA binding mixes. After 30min incubation at 30°C, binding reactions were loaded on 6% polyacrylamide gels and separated by electrophoresis in 0.5X TBE. Transfer to a nylon membrane and detection of biotin-labeled DNA were performed according to the manufacturer’s instructions (Thermo Scientific 20148 LightShift Chemiluminescent EMSA Kit).

### CUT&Tag experiment and data analysis

CUT&Tag were performed using endosperm at 10 DAP from *TaNF-YC6-RNAi-1* and Fielder plants. Two biological replicates were performed independently. The CUT&Tag experiment was performed exactly following protocols as previously described (Zhao et al., 2023). Finally, the purified PCR products were sequenced using an Illumina NovaSeq platform. For data analysis, the pipeline was largely based on the previous study (Zhang et al. 2023). The low-quality reads of of CUT&Tag library was filtered using fastp (v0.20.0) (Chen et al., 2018). the cleaned reads were mapped to wheat reference genome-Ref-Seq v1.1 (International Wheat Genome Sequencing, 2018) using bwa mem algorithm (0.7.17) (Li and Durbin, 2009). MACS2 (v2.1.2) was used for peak calling. For H3K27me3 peak calling, “--keep-dup all -g 14600000000 --broad --broad-cutoff 0.05” were used. The peaks were further annotated to the wheat genome using ChIPseeker (Yu et al., 2015). The whole genome was divided into three regions: promoter (−3000bp of TSS), genic (TSS to TES) and distal (other). The MAnorm package72 was used for the quantitative comparison of CUT&Tag signals between samples with the following criteria: abs (M value) > 1 and P < 0.05.

### CUT&Tag qPCR

CUT&Tag-qPCR was performed as previously described (Tian et al., 2023). Briefly, the DNA products of CUT&Tag were divided, 6µL was used as “Input”, and the other 18µL undergoing library PCR amplification and purification was used as “IP products.” Finally, the “Input” and “IP products” were dilute 30 times for qPCR assay. The primers for specific regions are provided in in Supplemental Data Set S8.

### Firefly luciferase complementation imaging (LCI) assay

The coding sequence of *TaNF-YB7-B-Hap1*, *TaNF-YB7-B-Hap2*, *TaNF-YC6-B* and *TaSWN-B-N* were coloned into pCAMBIA1300-35S-Cluc-RBS or pCAMBIA1300-35S-HA-Nluc-RBS vectors (Liu et al., 2018a) to generate fusion constructs. Four different vectors (e.g., nLUC-TaNF-YB7-B-Hap1, nLUC-TaNF-YB7-B-Hap2, cLUC-TaNF-YC6-B and cLUC-TaSWN-B-N) enabling testing of protein-protein interaction, were cotransfected into *N. benthamiana* leaves epidermal cells by Agrobacterium-mediated infiltration. After 3 days of incubation, the injected leaves were sprayed with 1 mM luciferin (Promega, E2940) and the LUC signal was captured using a cooled CCD imaging apparatus (Berthold, LB985). Each assay was repeated at least three times. Relevant primer sequences are given in Supplemental Data Set S8.

### Haplotype analysis of *TaNF-Y*

The single nucleotide polymorphisms (SNPs) and phenotypic data used for haplotype analysis were obtained from the publicly available database (https://opendata.earlham.ac.uk/wheat/under_license/toronto/WatSeq_2023-09-15_landrace_modern_Variation_Data/) (Cheng et al., 2023). The dataset contained genotype and partial phenotypic data for 827 Watkins landraces and 223 modern cultivars. The grain phenotype data (thousand-grain weight, grain protein content, and starch content) were collected at the John Innes Centre Field Experimental Station in 2014 (Supplemental Data Set S9). SNPs (MAF > 0.05) located in the gene upstream 2000bp, coding regions, and downstream 500bp were extracted and subjected to haplotype analysis using the R package “geneHapR” (v1.1.9) (Zhang et al., 2023).

### Statistics and data visualisation

For quantitative results, including three biological replicates and at least three technical replicates, the data is presented in the form of mean ± standard deviation. The means of two samples were compared using Student’s two-tailed *t* tests. Analysis of variance (one-way ANOVA) was conducted using default parameters in Graphpad Prism 8.0.2 software. Significant differences were determined by Student’s *t* test or one-way ANOVA: *, *P*<0.05 and **, *P*<0.01. Detailed statistical analysis data are shown in Supplemental Data Set S10.

## Data availability

Sequence data from this article can be found in the EMBL library (http://plants.ensembl.org/index.html) under the following accession numbers: *TaNF-YA3-D*, *TraesCS4D02G289600*; *TaNF-YB7-B*, *TraesCS7B02G121900*; *TaNF-YC6-B*, *TraesCS6B02G185700*; *TaAGPL1*, *TraesCS1A02G419600*; *TaGBSSII*, *TraesCS2A02G373600*; *TaAGPS1a*, *TraesCS7A02G287400*; *TaSSIIIa*, *TraesCS1A02G091500*; *TaSuS2*, *TraesCS2A02G168200*; *TaGli-α*, *TraesCS6B02G065856*; *TaGli-γ-700*, *TraesCS1D02G000700*; *TaLMW-400*, *TraesCS1D02G007400*; *TaNAC019-A*, *TraesCS3A02G077900*; *TaNAC019-B*, *TraesCS3B02G092800*, *TaNAC019-D*, *TraesCS3D02G078500*; and *TaSWN-B*, *TraesCS4B02G181400*. RNA-seq and CUT&Tag-seq data are available from the National Genomics Data Center (https://ngdc.cncb.ac.cn/?lang=zh) under accession number PRJCA022157.

## Supplemental data

**Supplemental Figure S1.** Phylogenetic and expression pattern of TaNF-Y components.

**Supplemental Figure S2.** Generation and characterisation of agronomic traits of TaNF-YA3-RNAi-1 and TaNF-YC6-RNAi-1.

**Supplemental Figure S3.** Grain starch granule pattern in TaNF-YA3-RNAi-1, TaNF-YC6-RNAi-1, and Fielder.

**Supplemental Figure S4.** Overview of RNA-seq data of TaNF-Y knock-down lines

**Supplemental Figure S5. T**he DNA binding capability and transcriptional activity of TaNF-Y components.

**Supplemental Figure S6.** TaNF-Y directly binds to the promoter and inhibits the expression of TaNAC019-B.

**Supplemental Figure S7.** The expression pattern of PRC2 components during endosperm development.

**Supplemental Figure S8.** TaNF-YC6 mediates the deposition of H3K27me3 at SSP coding genes.

**Supplemental Figure S9.** Starch granules pattern of *Tafie-C87* line.

**Supplemental Figure S10.** A proposed model for the direct and indirect regulation starch biosynthesis and SSP composition by TaNF-Y-PRC2 module.

**Supplemental Data Set S1.** The expression level of SSP and starch synthesis genes during wheat endosperm development.

**Supplemental Data Set S2.** List of differentially expressed genes between the *TaNF-YA3-RNAi-1*, *TaNF-YC6-RNAi-1* and Fielder.

**Supplemental Data Set S3.** The gene list with CCAAT motif in open promoter region of DEGs in *TaNF-Y RNAi* lines compared to Fielder.

**Supplemental Data Set S4.** List of differentially expressed genes between *Tanac019-cr* and Fielder.

**Supplemental Data Set S5.** List of genes with altered H3K27me3 levels in the *TaNF-YC6-RNAi-1* compared to KN199.

**Supplemental Data Set S6.** List of differentially expressed genes between *Tafie-C87* and KN199.

**Supplemental Data Set S7.** Association of *TaNF-YA/B/C* genes variation with grain development traits.

**Supplemental Data Set S8.** Primers used in this study.

**Supplemental Data Set S9.** Detailed information of the wheat accessions used in this research.

**Supplemental Data Set S10.** Detailed statistical analysis in this study.

## Author contributions

J.X. designed and supervised the research, J.X. and J.-C.C. wrote the manuscript. J.-C.C. did most of the experiments; H.-R.L did the GST pull down and EMSA assay; L.Z. Y.-J.L and H.Z. performed bio-informatics analysis; D.-Z. W. P.Z. did the haploid and selection analysis; X.-M. B. and X.-S.Z. provided *Tafie-C87* line; C.-F.Y. measured the protein content; Y.-Y.Y. provides *Tanac019-cr* transgenic wheats; X.-L.L., Y.-Y.Y., S.-B.X., X.-S.Z, and X-Y.Z. revised the manuscript; J.-C.C., L.Z., D.-Z.W., and J.X. prepared all the figures. All authors discussed the results and commented on the manuscript.

## Acknowledgements

We thank Professor Simon Griffiths (John Innes Centre, UK), Professor Cristobal Uauy (John Innes Centre, UK) and Professor Shifeng Cheng (Guangdong Laboratory for Lingnan Modern Agriculture) for sharing the Watkins genotypes and phenotypical data from preprint. We thank Yiman Yang (Nanjing Agricultural University) for help with figures layout.

---

**Supplemental Figure S1.**
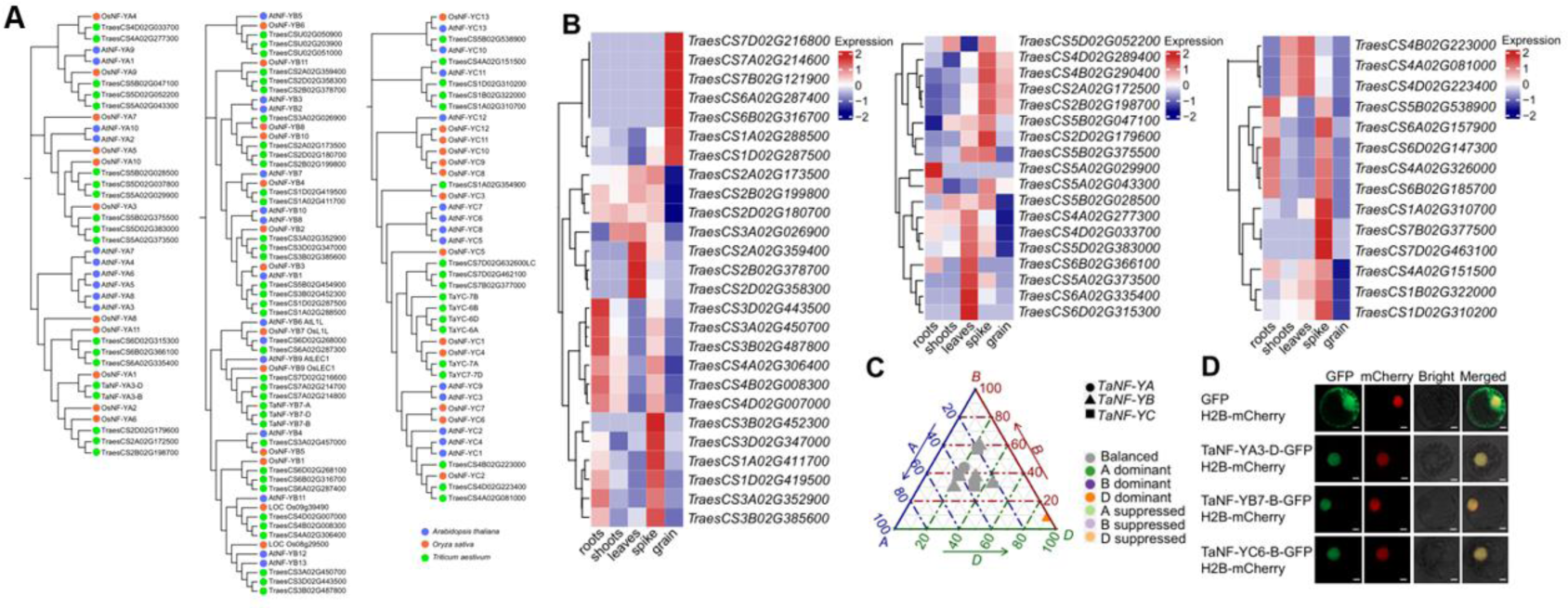
Phylogenetic and expression pattern of TaNF-Y components. A. Phylogenetic trees analysis of TaNF-Y proteins from different species. The TaNF-Y sequence from wheat, rice and *Arabidopsis thaliana* were used to construct the Phylogenetic tree. B. Heatmaps showing the expression level of TaNF-Y in different tissues of Chinese Spring, determined by RNA-seq. C. Homoeologs bias expression of TaNF-YA, TaNF-YB and TaNF-YC. D. Subcellular localization of TaNF-Y in wheat protoplasts. The free GFP and TaNF-YA3-D-GFP, TaNF-YB7-B-GFP and TaNF-YC6-B-GFP fusions under the control of the CaMV 35S promoter were transiently expressed in wheat protoplasts; 16 h after transformation, wheat protoplasts were observed using a confocal microscope. Scale bar, 0.5 µm.

**Supplemental Figure S2.**
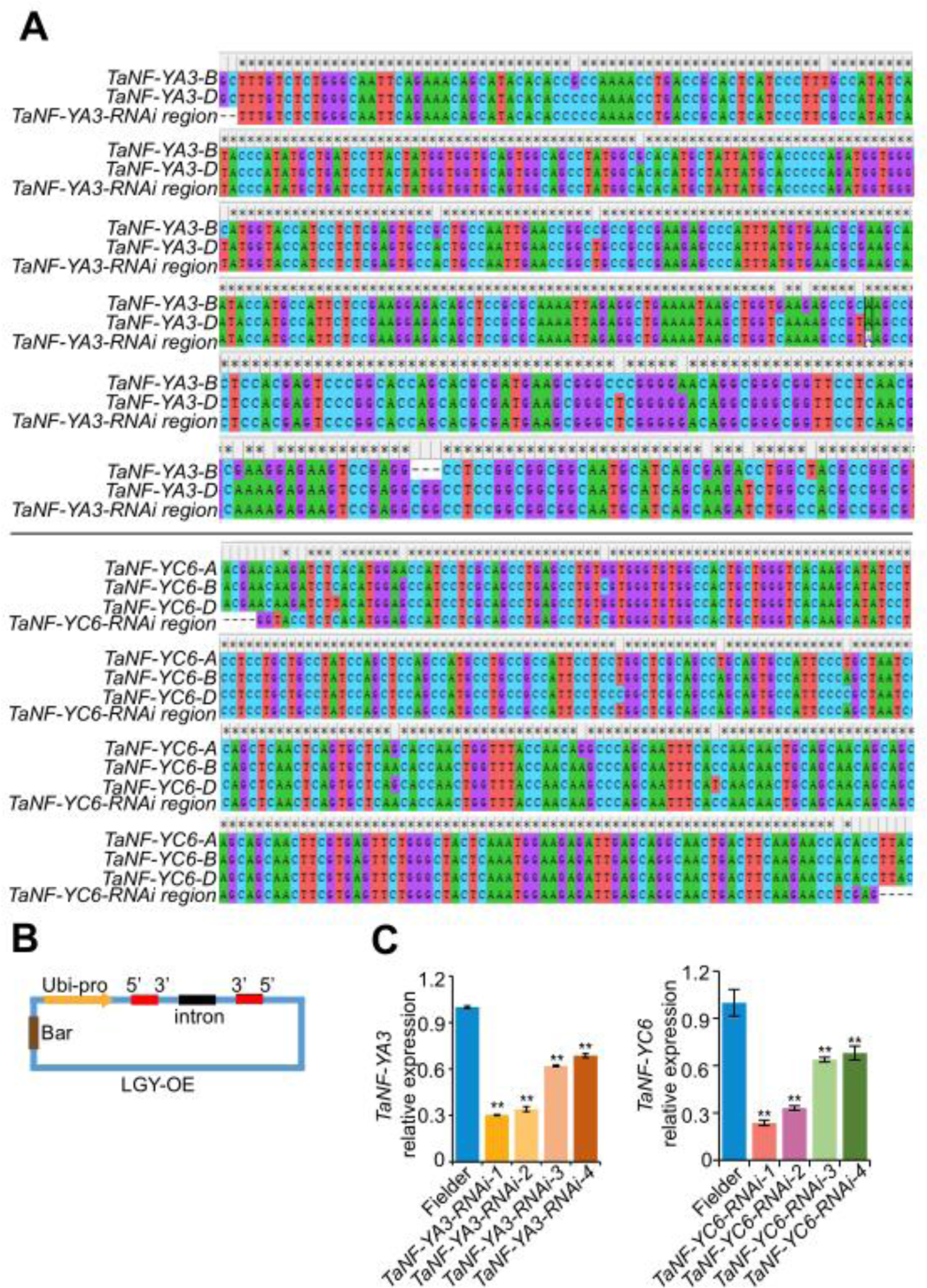
Generation and characterisation of agronomic traits of *TaNF-YA3-RNAi-1* and *TaNF-YC6-RNAi-1*. A. The DNA fragments of *TaNF-YA3* and *TaNF-YC6* for RNAi vector construction. B. Diagram showing the RNAi vector used in this study. C. Expression level of *TaNF-YA3*, and *TaNF-YC6* in knock down lines

**Supplemental Figure S3.**
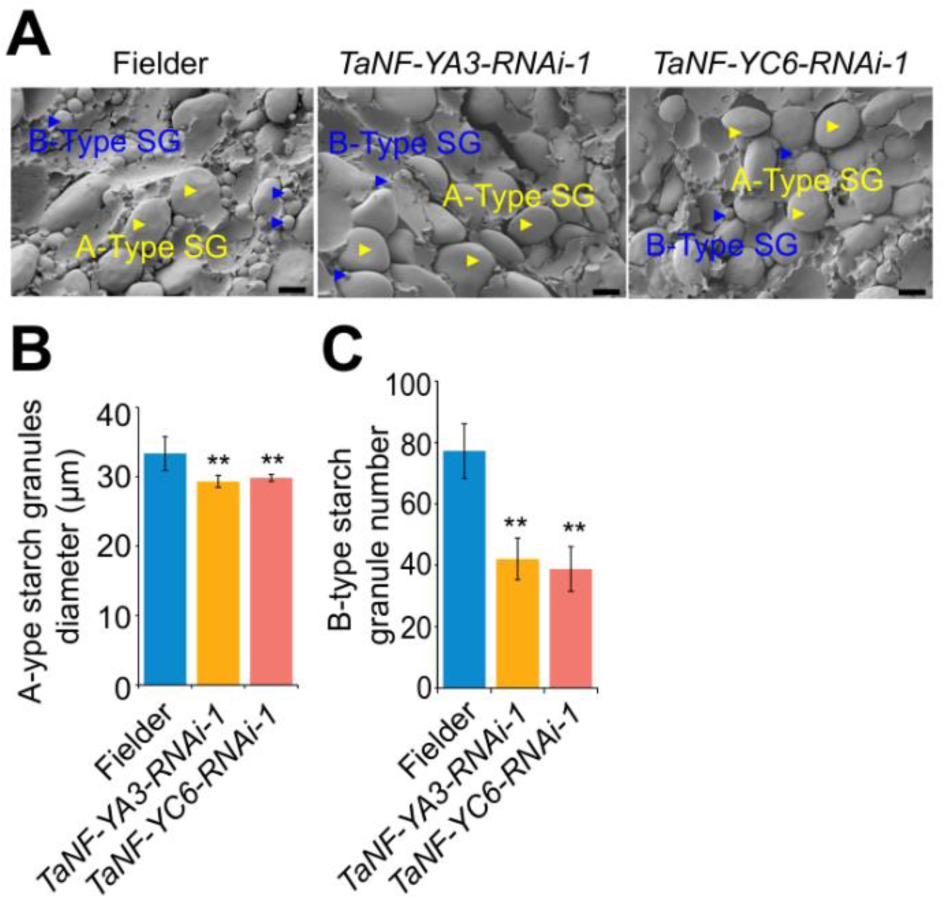
Grain starch granule pattern in *TaNF-YA3-RNAi-1*, *TaNF-YC6-RNAi-1*, and Fielder. A. Scanning electron microscopy of starch granules in the mature grain endosperm of Fielder, *TaNF-YA3* and *TaNF-YC6* knock-down lines. The A-type starch granules and B-type starch granules were indicated by blue and yellow triangles, respectively. Scale bar, 15 µm. B-C. statistical data of the diameter of A-type starch granules (B). the number of B-type starch granules (C). Statistical significance was determined by Student’s *t* test. *, *P* < 0.05; **, *P* < 0.01.

**Supplemental Figure S4.**
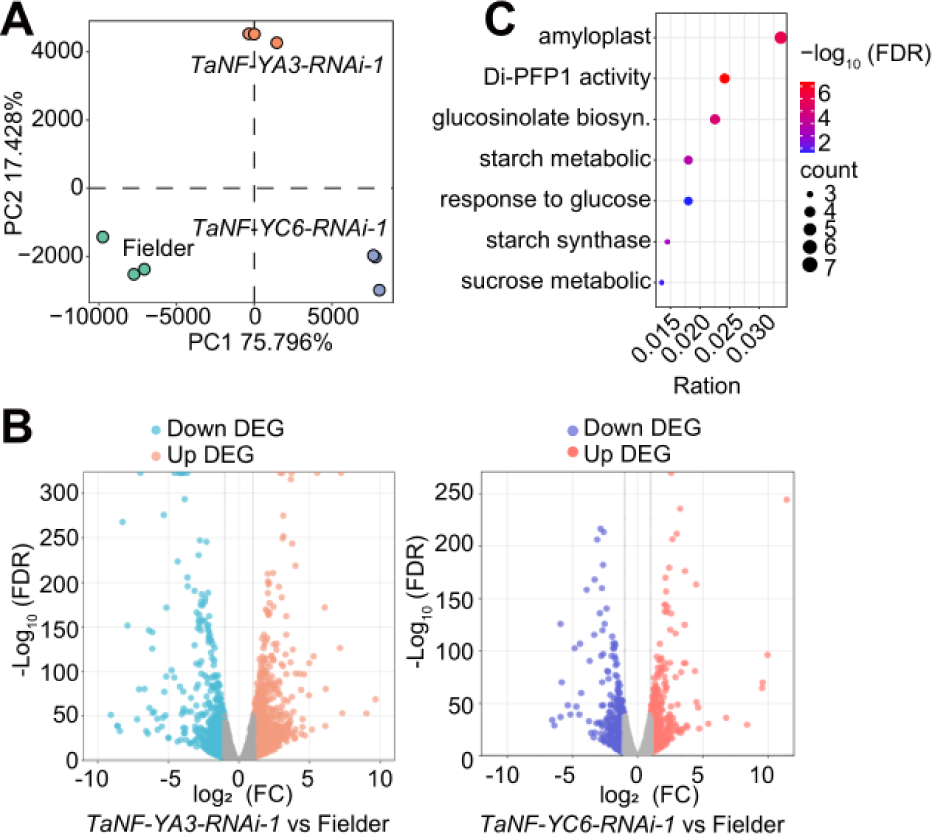
Overview of RNA-seq data of *TaNF-Y* knock-down lines. A. Principal component analysis (PCA) of transcriptome showing distinct lines. Each sample is represented by a dot, with three biological replicates sequenced for each line. B. Volcano plot showing the up- and down-regulated genes of *TaNF-YA3* and T*aNF-YC6* knock-down lines as compared to Fielder. C. Go enrichment analysis of shared down-regulated genes in the *TaNF-YA3-RNAi-1*, and *TaNF-YC6-RNAi-1*.

**Supplemental Figure S5.**
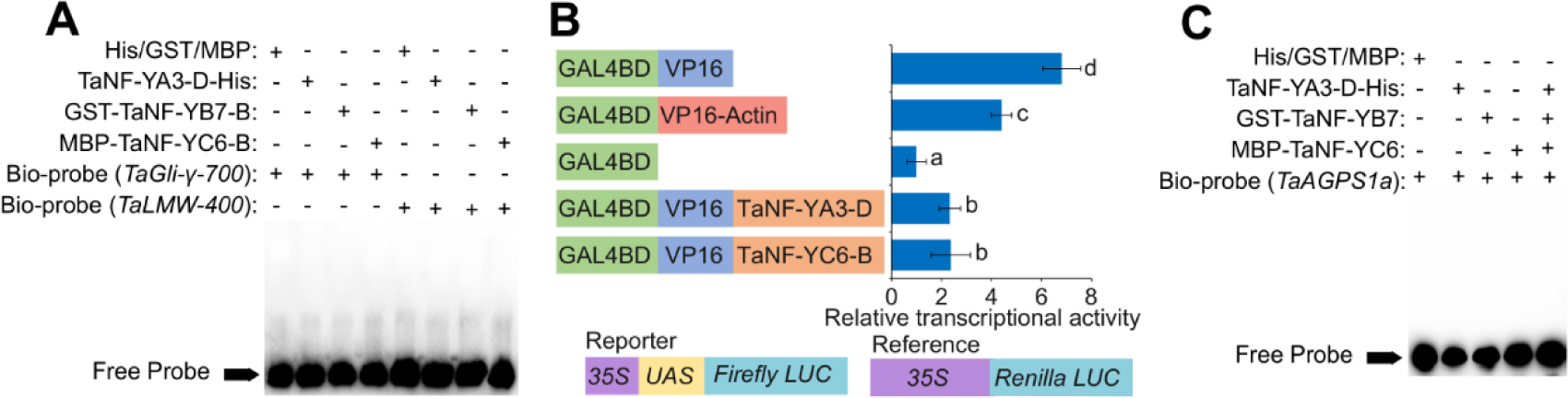
The DNA binding capability and transcriptional activity of TaNF-Y components. A. The EMSA assay show TaNF-YA3-D, TaNF-YB7-B, and TaNF-YC6-B cannot direct bind to the *TaGli-γ-700*, and *TaLMW-400* promoter. The recombinant protein of TaNF-YA3-His, GST-TaNF-YB7-B, and MBP-TaNF-YC6-B were used to assess the binding of these proteins to the *TaGli-γ-700* promoter. “+” and “-” indicate the presence and absence of the probe or protein, respectively. B. TaNF-YA3-D and TaNF-YC6-B show strong transcriptional inhibition activity in wheat protoplasts. Activities of firefly luciferase driven by the GAL4 binding element UPSTREAM ACTIVATION SEQUENCE (UAS) were measured. Renilla LUC was used as a reference and VP16, VP16-Fused-Actin were used as positive control, with mean ± SD of four biological replicates. Different letters indicate a statistically significant difference between different groups by one-way ANOVA and Tukey’s multiple comparison test at *P* < 0.05. C. The EMSA assay show TaNF-Y cannot bind to the *TaAGPS1a* promoter. The recombinant protein of TaNF-YA3-His, GST-TaNF-YB7-B, and MBP-TaNF-YC6-B were used to assess the binding of these proteins to the *TaAGPS1a* promoter. “+” and “-” indicate the presence and absence of the probe or protein, respectively.

**Supplemental Figure S6.**
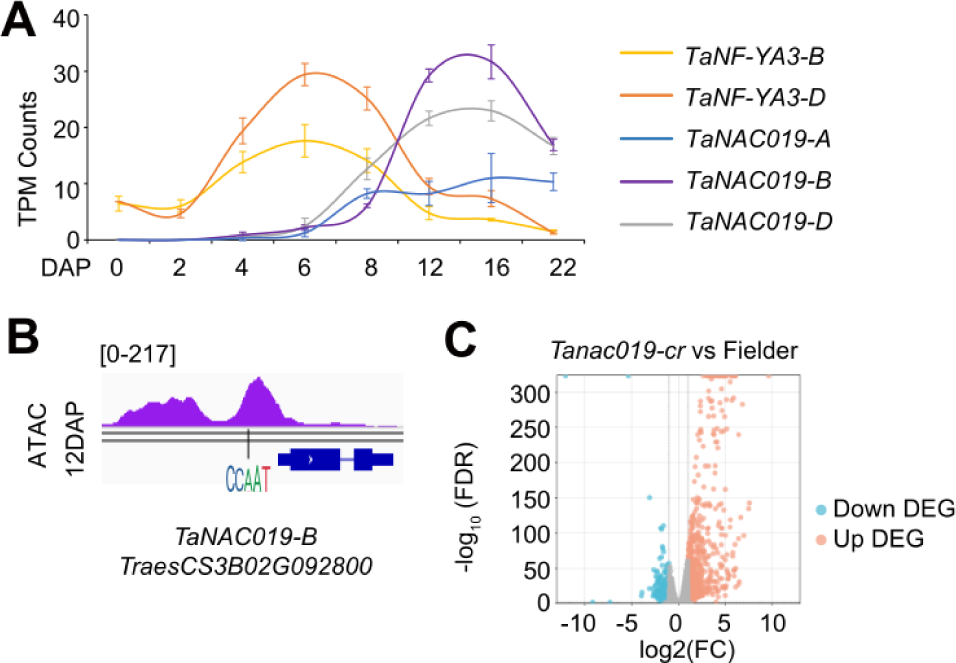
TaNF-Y directly binds to the promoter and inhibits the expression of *TaNAC019-B*. A. The alternating expression pattern of *TaNF-YA3* and *TaNAC019* during endosperm development. B. IGV shows the chromatin accessibility region of *TaNAC019-B* with a black vertical line indicating the CCAAT motif. C. Volcano plot showing the up- and down-regulated genes of the *Tanac019-cr* compare with Fielder.

**Supplemental Figure S7.**
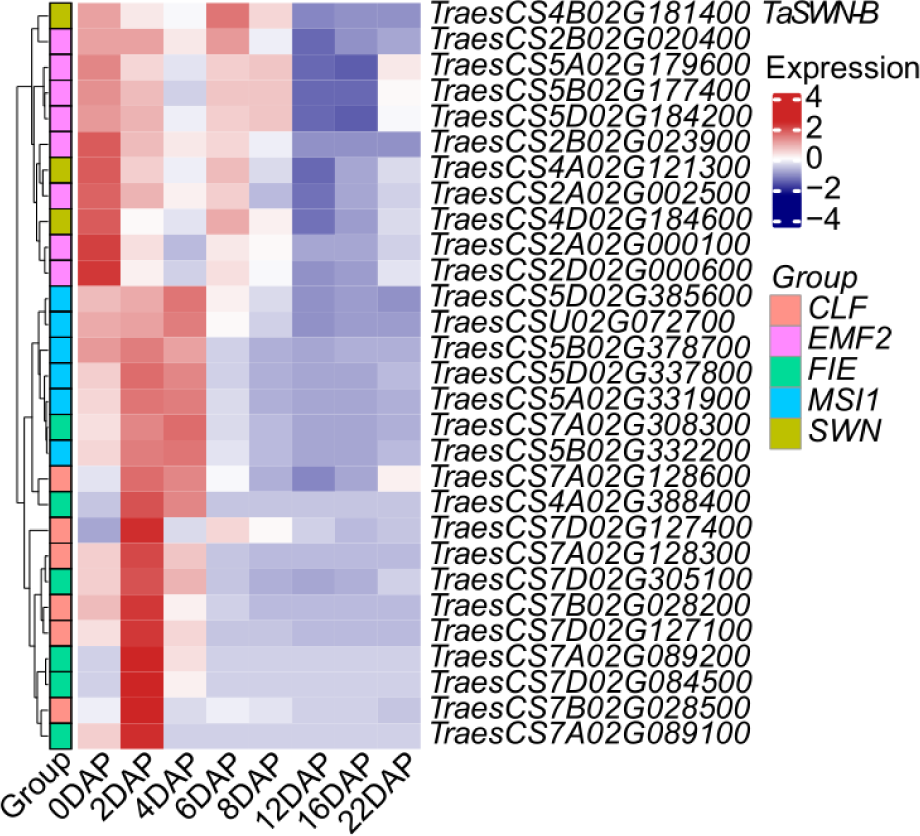
The expression pattern of PRC2 components during endosperm development. The expression pattern of PRC2 components during endosperm development.

**Supplemental Figure S8.**
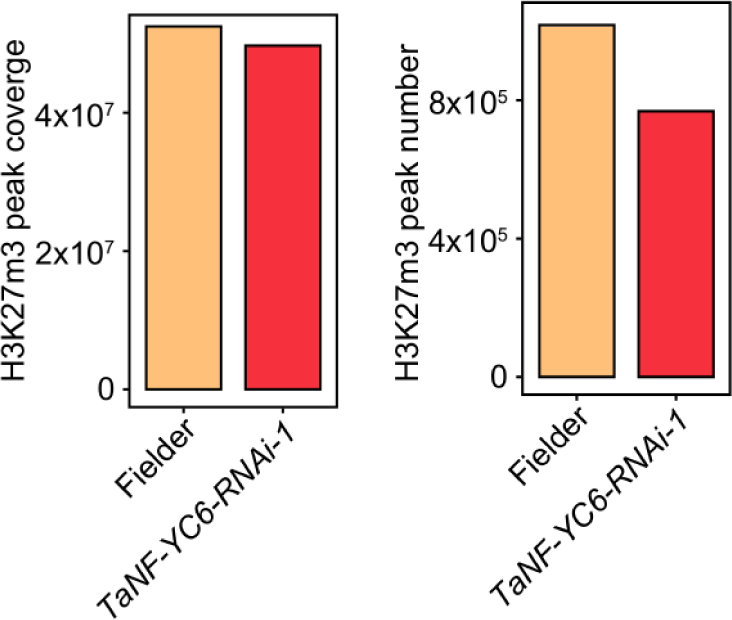
TaNF-YC6 mediates the deposition of H3K27me3 at SSP coding genes. Comparison of differences in peak coverage, and peak number of H3K27me3 modification on storage protein genes between *TaNF-YC6-RNAi-1* and Fielder.

**Supplemental Figure S9.**
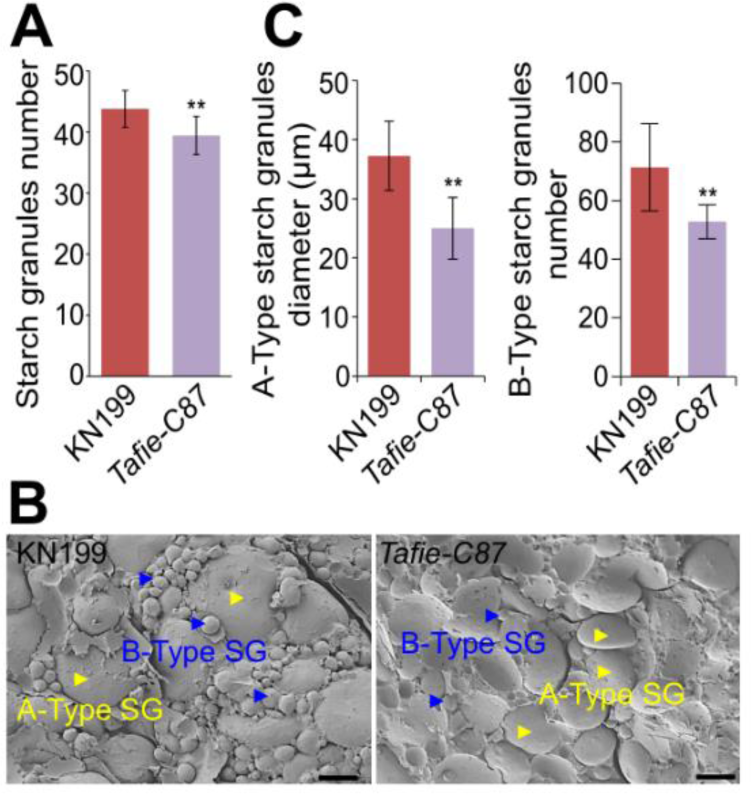
Starch granules pattern of *Tafie-C87* line. A. Statistics on the number of KN199 and *Tafie-C87* starch granules. Statistical significance was determined by Student’s *t* test. *, *P* < 0.05; **, *P* < 0.01. B-C. Scanning electron microscopy of KN199 and *Tafie-C87* endosperm starch (B). A-type starch granules diameter, and the number of B-type starch granules (C). The A-type starch granules and B-type starch granules were indicated by blue and yellow triangles in the scanning electron microscopy photos, respectively. Scale bar, 15 µm. Endosperm starch from ten plants was used to determine the diameter of complete A-type starch granules using Image J, with more than 60 starch granules measured for each plant. Statistical significance was determined by Student’s *t* test. *, *P* < 0.05; **, *P* < 0.01.

**Supplemental Figure S10.**
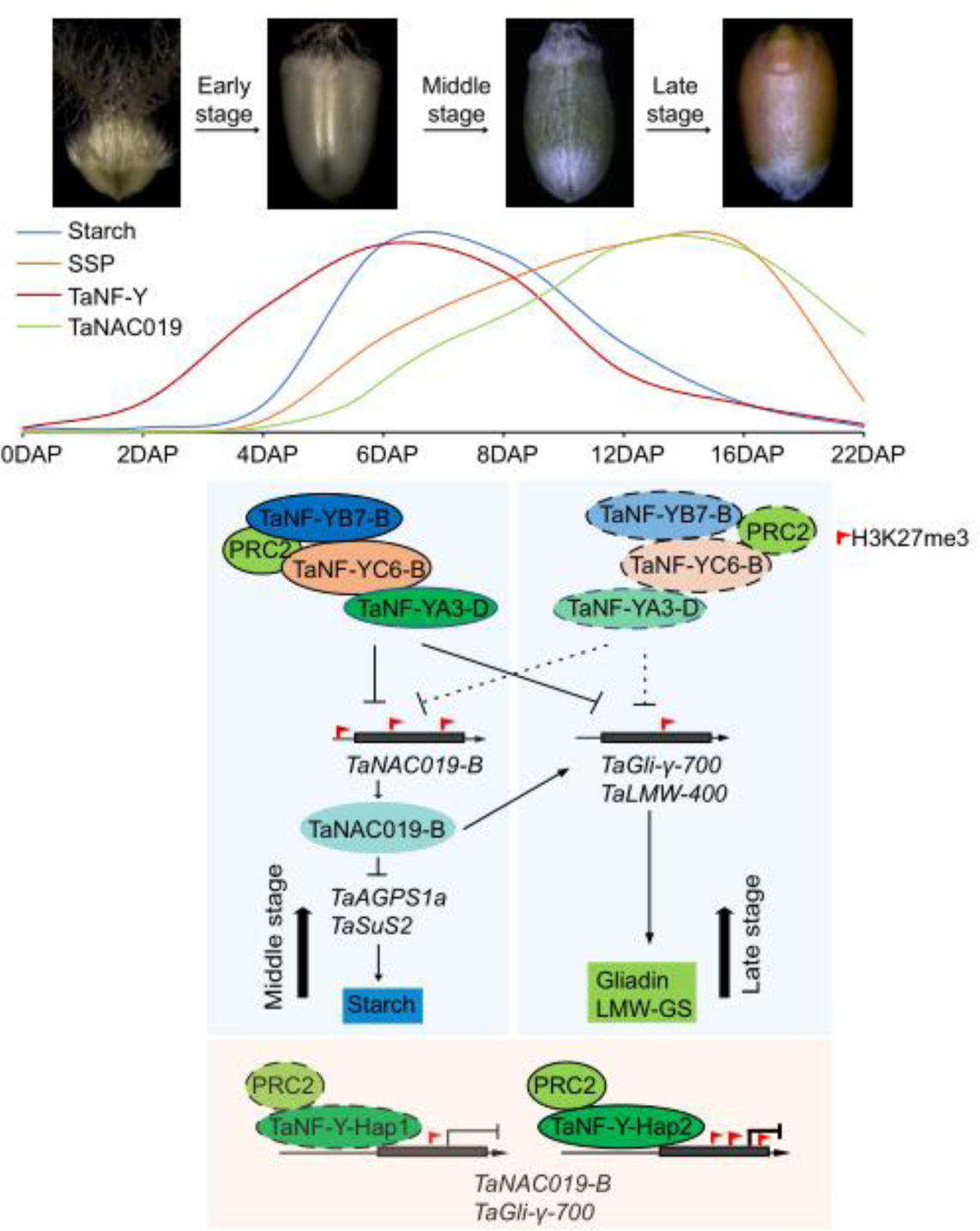
A proposed model for the direct and indirect regulation starch biosynthesis and SSP composition by TaNF-Y-PRC2 module. In the middle stage of wheat endosperm development (left), the higher expression of TaNF-YA3-D, TaNF-YB7-B, and TaNF-YC6-B form a complex with PRC2, recruited more K3K27me3 histone modification in *TaNAC019* and disinhibited the expression of some genes (such as *TaAGPS1a* and *TaSuS2*) involved in starch biosynthesis, thereby promoting the accumulation of starch. Simultaneously, higher modification of H3K27me3 at protein biosynthesis gene (such as *TaGli-γ-700*, *TaLMW-400*) suppresses their expression, therefore inhibiting the accumulation of storage protein. In the later stage of wheat endosperm development (right), the lower expression of TaNF-YA3-D, TaNF-YB7-B, and TaNF-YC6-B resulted in weakened interaction with PRC2, and alleviating transcriptional suppression of *TaNAC019* and protein synthesis genes, thus enhancing the accumulation of protein in endosperm while reducing the accumulation of starch. The TaNF-Y trimer complex containing the TaNF-YB7-B-Hap2 component has a higher interaction intensity with PRC2 and is more effective in inhibiting the transcription of downstream genes including *TaNAC019-B*, *TaGli-γ-700*, and, *TaLMW-400*.

